# Bacteroidales Secreted Antimicrobial Protein-1 receptor recognition is coupled to protease-activation to form bactericidal pores

**DOI:** 10.64898/2026.08.05.742979

**Authors:** S N Mostyn, K Flores, G Hedger, H Nagaraj, S L Rouse, L E Comstock, D Bubeck

## Abstract

Bacteroidales secreted antimicrobial proteins (BSAPs) are diffusible MACPF-domain toxins that mediate intra-species antagonism in the gut microbiota. Here we define the mechanism of action of BSAP-1 from *Bacteroides fragilis*, showing how target specificity encoded within the N- and C-terminal domains is coordinated with pore-forming activity of the MACPF. We show that specificity of the toxin for its receptor is mediated by an extended interface comprised of the BSAP-1 C-terminal domain and residues on the receptor that differ from the orthologous protein of BSAP-1 producing strains. On the surface of susceptible cells, BSAP-1 undergoes proteolytic cleavage of an N-terminal regulatory domain, triggering its assembly into oligomeric pores. Cryo-electron microscopy of membrane-inserted BSAP-1 reveals a 13-subunit transmembrane β-barrel pore formed through canonical MACPF rearrangements. Comparative modelling supports a conserved oligomerization mechanism across the BSAP family despite diversification of receptor-binding domains that target either proteins or glycan receptors. Together, these findings establish BSAP-1 as a receptor-targeted, protease-activated antibacterial MACPF toxin and provide a framework for understanding how gut Bacteroidales spatially restrict toxin activation to shape strain-level competition.

## Introduction

The order Bacteroidales comprises some of the most abundant and stable members of the healthy human gut microbiota [1]. Within this order, *Bacteroides fragilis* is notable in that it is frequently a symbiont in the human gut, but is also the anaerobic species most frequently isolated from extraintestinal infections, where it contributes to intraabdominal abscess formation and infections at extra-intestinal sites [2]. In addition, some *B. fragilis* strains produce the enterotoxin BFT [3], which has been linked to diarrhoeal disease [4] intestinal inflammation [5], and colorectal cancer [6, 7]. More broadly, high colonization levels of *B. fragilis* have been associated with diabetes [8] and mouse models suggest a possible link to Alzheimer’s disease [9]. Notably, *B. fragilis* strains differ substantially in their effects on both microbial community structure and host health, underscoring the importance of understanding how particular strains persist, compete, and exclude one another.

Gut Bacteroidales deploy multiple toxins to antagonize competitors, including those injected by contact-dependent Type VI secretion systems (reviewed [10]) and a repertoire of diffusible toxins [11–18]. Among the diffusible toxins are Bacteroidales secreted antimicrobial proteins (BSAPs), a family of toxins that contain a membrane attack complex/perforin (MACPF) domain and mediate intra-species killing [11–13]. While not all MACPF proteins have lytic activity [19, 20], several human immune proteins use MACPF domains to assemble pores that kill bacteria [21–23]. The cross-kingdom structural conservation of MACPF domains led to the hypothesis that the mechanism of BSAPs killing is through the formation of lytic pores in the outer membrane following interaction with the surface receptor [12, 24]. However, structural prediction tools fail to generate models consistent with a trans-membrane pore and neither oligomerization nor pores have been observed for BSAPs.

Previous studies showed that BSAPs target either outer membrane proteins [13, 24] or cell-surface glycans [12, 24] on susceptible strains. Because BSAPs are diffusible rather than contact-dependent, they must encode target specificity through receptor recognition. Gene-neighbourhood analyses suggest that BSAP genes co-evolve with adjacently encoded receptor variants [12, 24]. In producer strains, susceptible protein receptors are replaced by resistant orthologues [13, 24], whereas glycan-binding BSAPs are encoded adjacent to replacement glycosyltransferase genes that remodel the structure of lipopolysaccharide or lipooligosaccharide so it is no longer targeted by the BSAP [12, 24]. By replacing the BSAP receptor gene to encode an orthologue that no longer serves as a receptor, there is no need to encode an immunity gene to protect the producing cell from BSAP toxicity. These data suggest that BSAPs combine a conserved cytolytic core with receptor recognition, but the molecular basis of this selectivity remains poorly defined.

Several secreted antibacterial toxins of Bacteroidales are produced as inactive precursors that require processing by a surface protease of the clostripain family [16–18]. In *B. fragilis*, this clostripain is designated fragipain [25], a lipidated cysteine protease [26] initially characterized for its role in activating the disease-related enterotoxin BFT [3, 25, 27]. In *Phocaeicola dorei,* processing of antibacterial toxins BcpT and BSAP-3 with a related clostripain, DpnB, is necessary for toxicity [17]. However, how or if fragipain plays a role in controlling antibacterial activity of BSAPs of *B. fragilis* remains unknown.

Here we define a mechanism by which an antibacterial toxin achieves selective killing of closely related competitors in the gut microbiota. We show how BSAP-1 recognizes susceptible strains through a receptor-selective interaction mediated by its C-terminal domain and undergoes proteolytic activation to assemble transmembrane pores on target cells. Structural analysis of the membrane-inserted complex reveals that BSAP-1 forms a β-barrel pore through canonical MACPF rearrangements, while genetic, phenotypic, and biochemical data show that receptor engagement and target-cell processing are both required for bacterial killing. Together, these findings support a model in which receptor recognition, proteolytic processing, and pore formation are coordinated in a diffusible MACPF toxin deployed against competing bacterial cells.

## Results

### BSAP-1 C-terminal domain encodes target strain specificity

BSAP-1-mediated inter-strain competition within *B. fragilis* depends on selective recognition of the susceptible outer membrane receptor, referred to here as OMP-S, a β-barrel outer membrane protein predicted to transport hydrophobic molecules into the bacterium [24]. While a recent cryo-electron microscopy (cryo-EM) study established that BSAP-1 directly binds OMP-S, flexibility within the complex limited the map resolution and prevented molecular modelling of the interface [28]. BSAP-1 truncations showed that the C-terminal domain is necessary for bactericidal activity [28], but how BSAP-1 distinguishes the susceptible receptor OMP-S from the resistant orthologue (OMP-R) on producing strains, despite sharing 69.8% sequence identity, remained unknown. To address this, we modelled the BSAP-1-OMP-S complex using Alphafold3 (AF3) [29] (Fig. 1a). Although residues within the BSAP-1 C-terminal domain were predicted with a lower confidence than the central MACPF domain, the local distance difference test (pLDDT) scores of these residues increased when in complex with OMP-S and decreased within OMP-R (Supplementary Fig. 1a, Supplementary Fig. 1a, and Supplementary Table 1). Further analysis of the interface pair-wise alignment error (iPAE) highlights several contiguous residues within the BSAP-1 C-terminus that are within 5 Å from OMP-S and are consistent with a confident interface prediction (Fig. 1a, Supplementary Table 1). By contrast, only three residues satisfied these criteria for the OMP-R-BSAP-1 model, suggesting a low confidence prediction (Supplementary Fig. 1c). No other notable interactions were identified between OMP-S and BSAP-1. AF3 modelling suggests that the BSAP-1 C-terminal domain wraps around a groove formed by the extracellular loops of OMP-S (Fig. 1a), which are notably longer than those present in OMP-R (Supplementary Fig. 1d). The most extensive interaction interface comprises an inter-molecular β-sheet stabilized by a network of hydrogen bonds formed across back-bone atoms (S356-V359) of BSAP-1 and a β-hairpin extension of OMP-S lacking in the resistant orthologue (Fig. 1a and Supplementary Fig. 1d).

**Figure 1.**
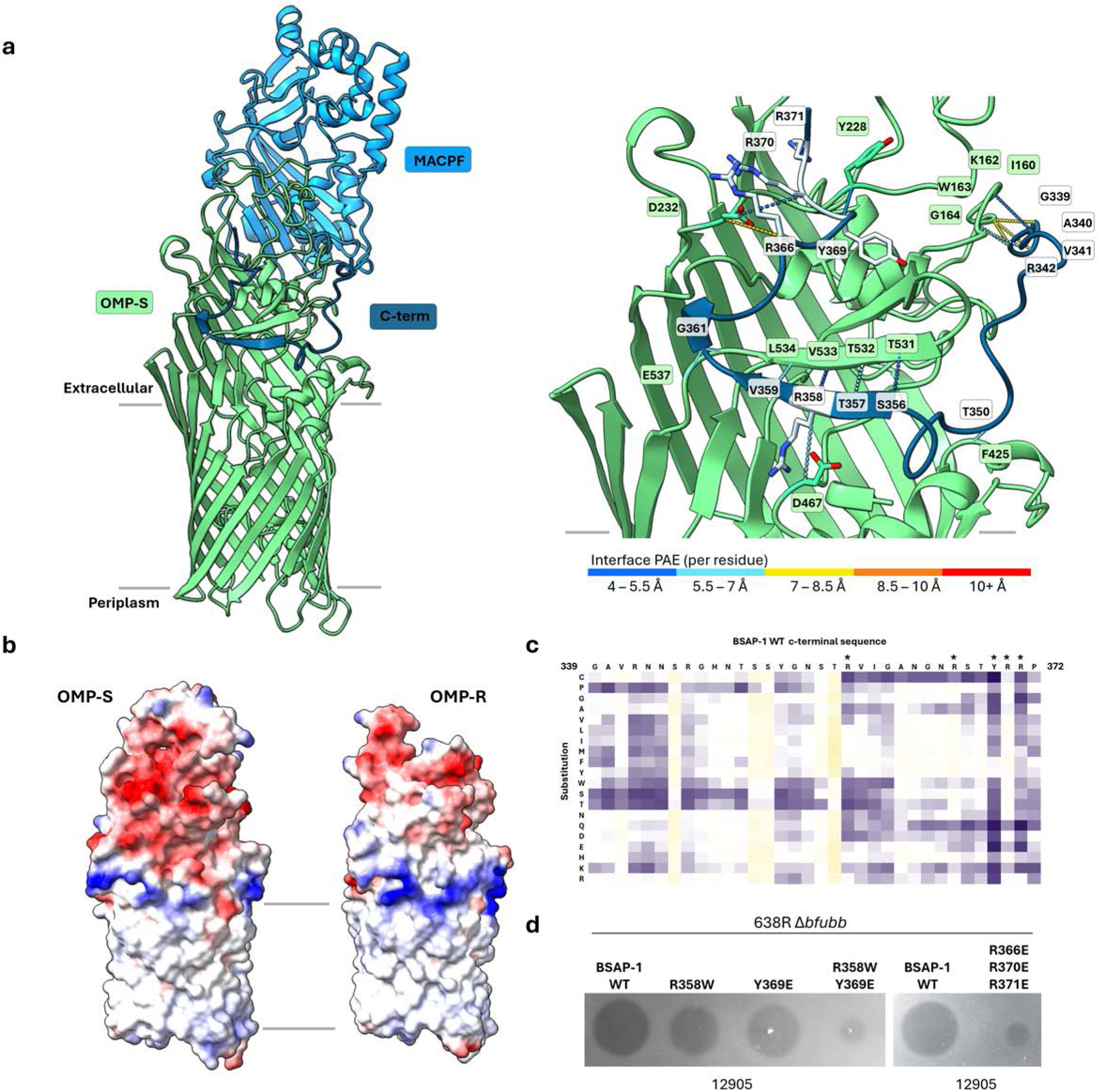
Molecular basis of OMP-S recognition by BSAP-1. **(a**) AF3 predicted binding interface between BSAP-1 (blue) and OMP-S (green). The predicted periplasmic, outer membrane spanning, and extracellular regions of OMP-S are shown. Inset: Zoom in of the predicted interface showing the per residue predicted aligned error (PAE) as colored dotted lines. Interface residues are defined as having Cα distance cutoffs: 5 Å for backbone interactions, and 8 Å for salt bridges. Full list of interactions in Supplementary Table 1. Sidechains of BSAP-1 residues mutated in the *B. fragilis* 638R strain and used in antagonism assays are shown (grey). Residues on OMP-S that are within the specified interface cutoff distances of mutated residues are shown (sidechains, green). (**b**) Coulombic electrostatic potential ranging from −10 (red) to 10 (blue) kcal/(mol·e) calculated from the OMP-S and OMP-R AF3 models, highlighting the differences in the extracellular negatively charged patch. (**c**) Far-Western analysis of BSAP-1 C-terminal peptide array shows interacting residues. BSAP-1 C-terminal 33-mer peptide (listed across the top) having all possible amino acid substitutions (listed down the side) was constructed on a peptide array. The ability of each peptide to interact with OMP-S was assessed by far-Western blotting, using a histidine-tagged purified OMP-S and detection using a histidine tag antibody. Signals for each peptide spot were quantified and normalised for the WT spot in each column. Purple: loss of signal, white: no change, yellow: gain of signal. Residues chosen for mutational analyses in *B. fragilis* 638R indicted by asterisks. Raw data shown in Supplementary Fig. 1b and Supplementary Table 2. (**d**) Agar spot overlay assay showing killing of *B. fragilis* 12905 by *B. fragilis* 638R expressing BSAP-1 mutants. Zone of clearing (dark circles) indicate that the spotted *B. fragilis* 638R strain listed at the top prevents *B. fragilis* 12905 growth in the overlay.

Additionally, several C-terminal domain arginine residues map on to a negatively charged surface of OMP-S that is lacking in OMP-R (Fig. 1b). The AF3 model of the interface is consistent with the cryo-EM density of the complex (Supplementary Fig. 2), supporting the hypothesis that BSAP-1 C-terminal domain binds extracellular loops of OMP-S through an extended interface. To validate the predicted interface, we performed a far-western blot analysis of a peptide array based on the C-terminal BSAP-1 residues (Fig. 1c, Supplementary Figure 1b, Supplementary Table 2). Substitution to a range of amino acids at positions R358 and Y369 reduced OMP-S detection. Besides the cysteine substitution, R358W and Y369E has some of the largest effects on binding in the peptide array. To test if these alterations reduced BSAP-1 mediated killing in the context of the bacterial cell, these substitutions were made in the *B. fragilis* 638R Δ*bfubb* strain to prevent killing by the Bfubb toxin [14], and tested for killing activity against sensitive strain *B. fragilis* 12905 (Fig. 1d). Although the individual point mutations had minimal effects as assessed in the spot agar overlay assay, the combined R358W/Y369E substitution showed substantially reduced killing activity. Because the AF3 predicted interface pairs an arginine-rich BSAP-1 C terminus with a negatively charged region unique to OMP-S (Fig. 1 and Supplementary Fig. 1d), we next tested whether activity depended on the combined contribution of multiple charged residues. Substitution of the three terminal arginine residues reduced OMP-S binding in the peptide array to varying degrees (Fig. 1c, Supplementary Fig. 1b, and Supplementary Table 2). Indeed, a triple substitution of these R366E/R370E/R371E showed markedly reduced killing activity in overlay assays (Fig. 1d), in agreement with the seven-residue C-terminal truncation eliminating activity [28]. Together, these data show that BSAP-1 distinguishes susceptible and resistant OMP orthologues through a distributed C-terminal interaction surface composed of electrostatic and hydrogen-bonding contacts.

Although the AF3 predicted models converged on a common C-terminal binding interface, the MACPF domain of BSAP-1 adopted multiple orientations relative to OMP-S, consistent with the marked flexibility observed previously for the detergent-solubilized receptor-bound complex [28]. We therefore used atomistic molecular dynamics simulations to test whether a membrane environment stabilizes a preferred orientation of BSAP-1 on receptor-bound OMP-S (Fig. 2). Since the N-terminal domain is predicted to be disordered, we used AF3 to model the complex using only the C-terminal and MACPF domains of BSAP-1. Throughout the simulation, the BSAP-1 C-terminal domain remained stably associated with OMP-S (Fig. 2a, b), further supporting the robustness of the predicted model. Superposition of the C-terminal residues from evenly sampled frames across the trajectory showed minimal variation (Fig. 2b) and although the MACPF domain remained stably folded (Supplementary Fig. 3b), it retained substantial mobility relative to the receptor (Supplementary Fig. 3a). Flexibility between the two domains was enabled in part by a pair of glycine residues linking the C-terminal domain to the MACPF (Fig. 2c, d). Analysis of protein-lipid contacts showed the expected high lipid occupancy around the transmembrane belt of the OMP-S β-barrel, whereas residues within the BSAP-1 MACPF domain exhibited only transient membrane contacts over the course of the simulation (Fig. 2e, Supplementary Fig. 3c). Thus, receptor engagement does not rigidly orient the MACPF domain or trigger further conformational changes in this domain, even in a membrane context. Because these simulations were performed in a simple DOPC bilayer, additional features of the native *B. fragilis* outer membrane may further influence MACPF domain orientation at the target surface. Simulations also lacked the disordered N-terminal domain, which may also play a role in directing BSAP-1 activity.

**Figure 2.**
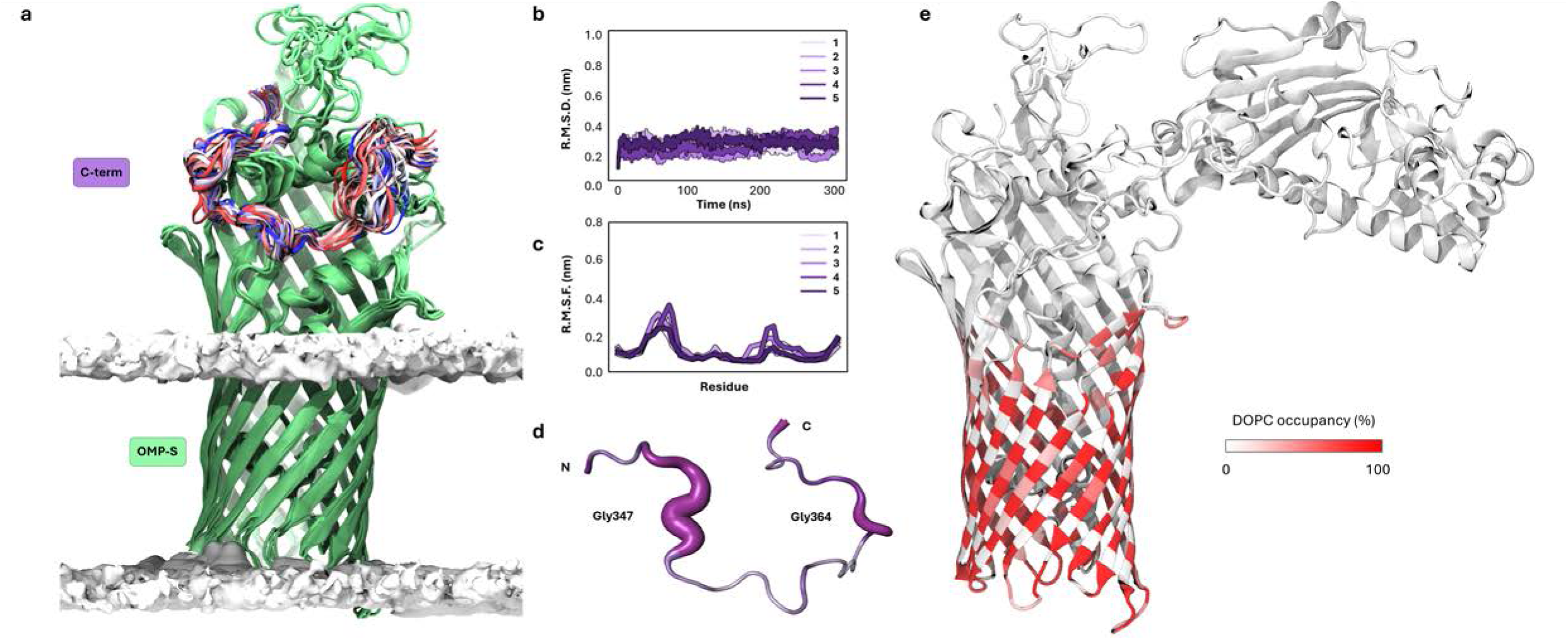
Molecular dynamics simulation reveals flexibility of BSAP-1-OMP-S complex in a membrane environment. **(a)** Overlay of 50 evenly distributed structures of the C-terminal domain of BSAP-1 over a combined 5 x 300 ns ensemble. The structures are coloured red-white-blue according to simulation time. BSAP-1 MACPF is hidden, and visible in Supplementary Fig. 3a. The phosphate region of the lipid bilayer is shown as a white surface. OMP-S is shown as green ribbons. (**b)** Root mean square deviation (RMSD) of the C-terminal domain in complex with OMP-S. (**c-d)** Root mean square fluctuation (RMSF) of each residue in the C-terminal domain. Flexible residues are visualised: stable (thin), flexible (thick). N and C termini of the BSAP-1 interacting residues are indicated. (**e)** Lipid occupancy of the complex averaged across simulations. Raw data shown in Supplementary Fig. 3c for all residues.

### N-terminal processing of BSAP-1 allows for controlled release of pore-forming residues

Clostripain-family surface proteases activate diverse extracellular toxins in Bacteroidales species [16–18, 25], raising the possibility that fragipain also regulates BSAP-1 activity. Inspection of the BSAP-1 sequence identified a single candidate fragipain cleavage site within the N-terminal region that is conserved across BSAP family members (Fig. 3a). To test whether cleavage at this site is required for activity, we introduced an R38N substitution predicted to disrupt protease cleavage and assessed antagonistic activity of a *B. fragilis* 638R producer strain expressing the variant (Fig. 3b). This substitution abolished BSAP-1-mediated killing of susceptible strains, suggesting that cleavage at R38 is required for antagonism. To distinguish whether BSAP-1 is cleaved by the producing strain during secretion or the target strain upon binding, we deleted *fpn* from producing strain 638R and sensitive strain CL03T12C07, a *B. fragilis* strain in which deletion mutants can be constructed, and assessed BSAP-1 activity. Deletion of *fpn* in the producing strain had no effect on susceptible strain killing. By contrast, purified BSAP-1 failed to kill *B. fragilis* CL03T12C07 deleted for *fpn*. These results indicate that BSAP-1 cleavage by the producing cell is not necessary for activation. Instead, cleavage at the surface of the sensitive cell enables killing.

**Figure. 3.**
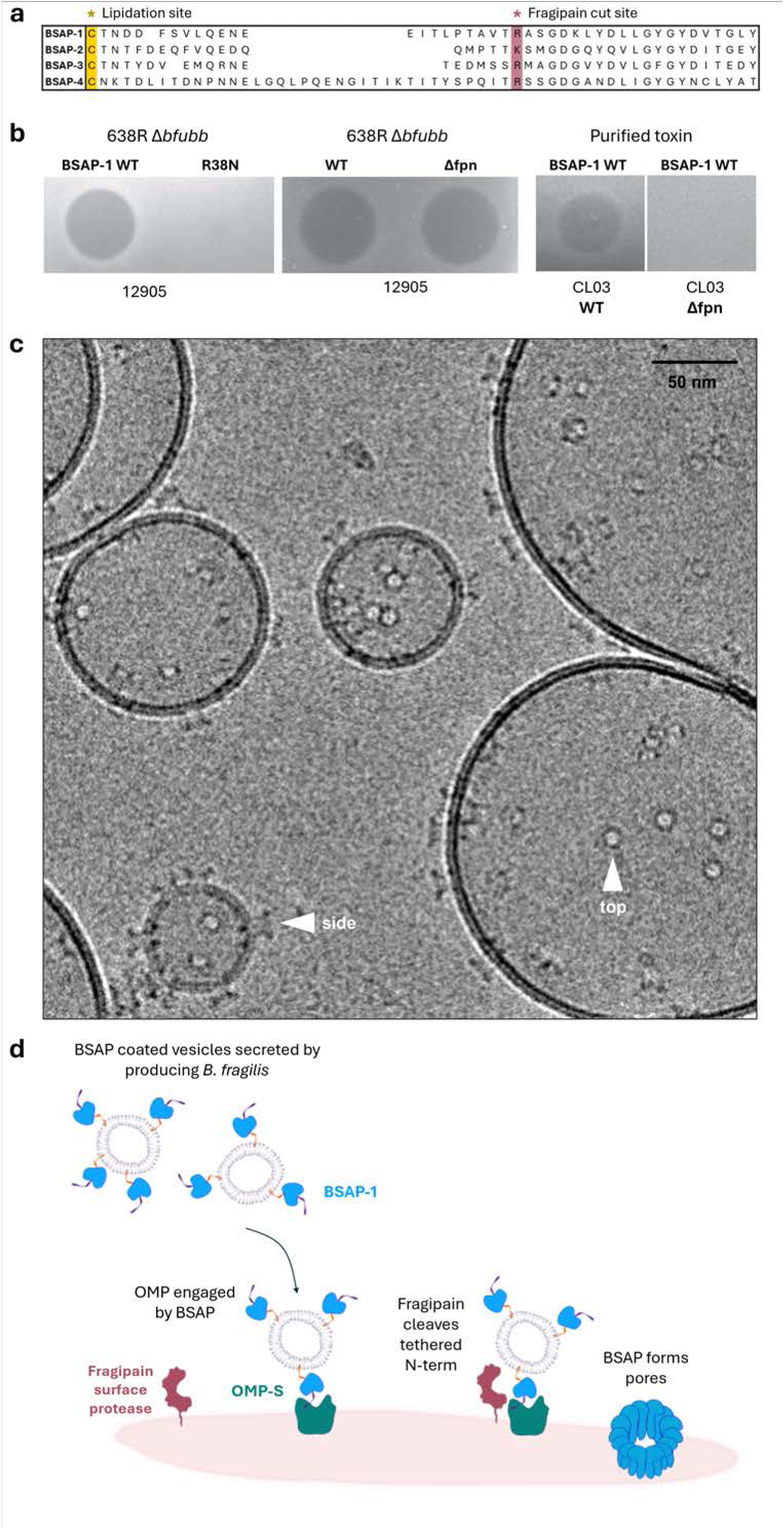
Proteolytic activation of BSAP-1 triggers pore formation. **(a)** BSAP family N-terminus sequence alignment. Predicted lipidated residue (yellow) and fragipan cut site (pink) are indicated. (**b)** Spot agar overlay assay assessing killing of BSAP-1 susceptible *B. fragilis* strains. Growth inhibition of *B. fragilis* 12905 by 638R Δ*bfubb* wildtype BSAP-1 or R38N BSAP-1 variant left panel). Growth inhibition of *B. fragilis* 12905 by 638RΔ*bfubb* or 638R Δ*bfubb*Δ*fpn* (middle panel). Growth inhibition of the genetically manipulable BSAP-1 susceptible CL03 or the fragipain deletion strain CL03 Δ*fpn* by purified wildtype BSAP-1 (right panel). (**c)** Representative cryo-EM image showing trypsin digest of BSAP-1 produces pores in liposomes. Top and side views of the pore are indicated with white arrows. (**d)** Proposed activation mechanism for formation of BSAP-1 pores in target membranes.

To determine whether proteolytic cleavage is sufficient to trigger pore formation, we treated BSAP-1 with trypsin, which cleaves at the same site as the clostripain, in the presence of liposomes and visualised complexes by cryo-EM (Fig. 3c). In the absence of receptor, digestion of BSAP-1 resulted in oligomerization of the toxin and formation of transmembrane pores. Together, these data support a model in which BSAP-1 is proteolytically processed at its conserved N-terminal cleavage site to direct pore formation on the surface of susceptible strains (Fig. 3d).

BSAPs are produced as surface-tethered outer membrane lipoproteins, with the lipid linked to the N-terminal cysteine residue following signal peptide removal (Fig. 3a). It has been shown that lipidation is critical to BSAP-1 mediated killing [28], likely because it allows sorting of the protein to the outer membrane. In addition to triggering conformational changes required for pore formation, cleavage at R38 likely disengages BSAP-1 from outer membrane vesicles (OMVs) and frees the MACPF to penetrate target membranes. Taken together, proteolytic activation of OMV delivered BSAP-1 provides a second checkpoint after receptor recognition to spatially restrict toxicity.

### Cryo-EM structure of the oligomeric BSAP-1 pore

To understand the structural transitions of BSAP-1 upon proteolytic activation, we solved the cryo-EM structure of the BSAP-1 pore inserted into lipid bilayers (Fig. 4, Supplementary Fig. 4). Two-dimensional class averages revealed a highly homogeneous complex comprised of thirteen BSAP-1 monomers. Applying C13 symmetry, we calculated a 2.7 Å reconstruction and built an atomic model of the transmembrane pore (Fig. 4, Supplementary Fig. 4, Supplementary Fig. 5, and Supplementary Table 3). Our structure shows that the MACPF domain forms a giant β-barrel pore with a lumen of 60 Å and height of 120 Å (Fig. 4a). In the absence of OMP-S, the C-terminal receptor-binding domain was disordered, and no density was observed for the cleaved N-terminal residues. A comparison with the predicted soluble state revealed that BSAP-1 pore formation is accompanied by displacement of the β5 strand of the central MACPF domain (Fig. 4b). This transition exposes the oligomerization interface in a manner analogous to the transition described for CDC [30, 31] and CDC-like toxins [32]. In the assembled pore, these displaced residues adopt an extended helix-turn-coil conformation that lines the lumen and contributes to stabilization of the oligomer. Although this motif is conserved across MACPF containing proteins [32–34], it is significantly longer in BSAP-1 and makes extensive contacts with the neighbouring monomer (Fig. 4c). This arrangement is supported by a highly conserved tryptophan residue that packs against the core β-barrel (Fig. 4b), consistent with previous functional data showing that mutation of the corresponding residue abolishes BSAP-1 activity [11].

**Figure 4.**
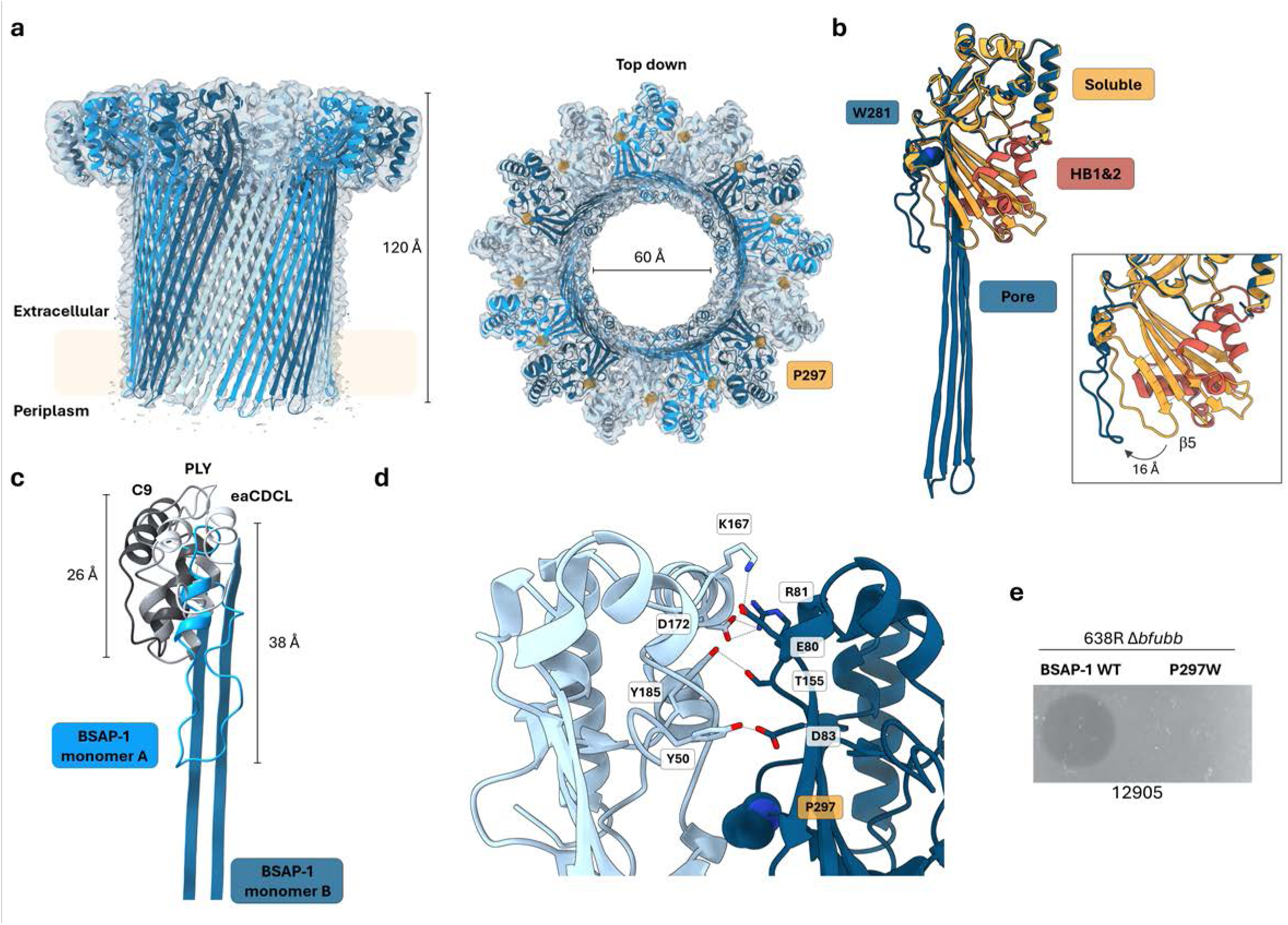
Cryo-EM structure of the BSAP-1 pore. **(a)** Cryo-EM reconstruction of the BSAP-1 pore (transparent surface) overlayed with a molecular model for the 13-mer (monomers alternating shades of blue). P297 (orange spheres) forms a key interaction at the oligomerization interface. (**b)** Superposition of the BSAP-1 pore structure with an AF3 prediction for a soluble monomeric conformation shows the transition of two helical bundles (HB1 and HB2) into transmembrane β-hairpins. Pore formation is accompanied by the unbending of the central MACPF domain and the displacement of the MACPF β5 strand, which forms the helix-turn-helix motif that lines the interior of the β-barrel (inset). This new conformation is stabilized in part by the aromatic sidechain of W281 (highlighted as blue spheres), a residue shown to be essential for BSAP-1 activity [11]. (**c**) Superposition of the pore conformations of BSAP-1, Complement component 9 (C9, PDB: 6H04), pneumolysin (PLY, PDB: 5LY6), and a CDC-like toxin (eaCDCL, PDB: 9CCP) shows that the helix-turn-helix motif of BSAP-1 is significantly longer and latches across the neighbouring monomer during oligomerization. **(d)** BSAP-1 oligomerisation interface residues indicated as sticks with h-bonds shown as dotted lines. P297 is highlighted as spheres. N-terminal loop is hidden for clarity. (**e)** Spot agar overlay assay showing 638RΔ*bfubb* BSAP-1 P297W variant loses killing activity of the susceptible *B. fragilis* strain 12905.

Displacement of β5 together with unbending of the central MACPF allows polymerization of BSAP-1 monomers (Fig. 4). At the oligomer interface, a proline residue from one monomer packs against a neighbouring protomer and may help define the pore geometry (Fig. 4a, d). Substitution of this residue with tryptophan (P287W) abolished BSAP-1 activity (Fig. 4e), supporting the conclusion that oligomeric pore formation is required for inter-strain antagonism.

To test whether this extends across the wider BSAPs, we generated AF-based pore models for BSAP-2, BSAP-3, and BSAP-4 using the BSAP-1 pore as a template. Although the corresponding proline is not conserved in primary sequence (Supplementary Fig. 6), each model contained an analogous proline located at the oligomer interface (Supplementary Fig. 7). Intriguingly, comparative analysis of BSAP pore models showed that glycan-binding BSAPs, including BSAP-2 and BSAP-3, contain a longer predicted transmembrane hairpin than protein binding BSAPs (Supplementary Fig. 6 and Supplementary Fig. 7). Alignment of these models relative to the membrane suggest the longer β-hairpin of glycan binding-BSAPs extends further into the periplasm, which may be relevant for their mechanism of action.

## Discussion

Our current understanding of inter-strain antagonism mediated by diffusible toxins of *B. fragilis* is based largely on genetic and phenotypic analyses that have identified bactericidal toxins and receptors on target strains [12, 24]. However, the molecular details for how this specificity is achieved and the mechanism by which BSAP-1 kills bacteria remains poorly understood. Here, we combined AF3 modelling with mutagenesis assays to reveal residues within the C-terminal domain of BSAP-1 that are essential for recognition. In addition, our cryo-EM structure of BSAP-1 pores in model membranes together with bacterial genetic and phenotypic assays, support a model in which fragipain cleaves BSAP-1 to trigger lytic pores on target cells. Together, our work provides a series of molecular snapshots of BSAP-1 activation and shed new light on how a diffusible antimicrobial toxin precisely targets competing *B. fragilis* strains in the gut microbiota.

Target specificity is essential to prevent bystander damage of pore-forming proteins. Cholesterol-dependent cytolysins (CDCs) are bacterial pore-forming toxins that encode selectivity for eukaryotic targets through a lipid specificity [34, 35]. Similarly, the human immune system deploys perforin-2 on bacterial surfaces or damaged human cells by binding negatively charged lipids [36]. Prior work established that BSAP-1-mediated bacterial antagonism depends on protein receptor recognition by the C-terminal domain [28], instead of lipids to direct activity. Our mutagenesis data now provide a molecular blueprint for engineering BSAP-1 specificity, showing that receptor recognition requires a combination of electrostatic and hydrogen bonding interactions with the receptor. Such a dependency across the broader C-terminal domain provides a selectivity that is robust against evolutionary pressure escape mutants. This principle likely extends to other BSAPs that couple a conserved MACPF pore-forming domain to a C-terminal domain used to discriminate target strains through cell-surface glycans [12].

BSAPs bind target receptors on rival bacterial strains at varying distances in the gut, ranging from proximal local niches to more distal spatial ranges. *B. fragilis* partially overcomes this spatial challenge by deploying OMVs, which contain proteins [37, 38] including BSAPs [11], polysaccharides [39] and small RNAs [40]. Tethering of BSAPs to OMVs may allow for delivery of a critical concentration of BSAP monomers to the target cell. Upon arrival BSAPs need to traverse flip from the OMV membrane to the target membrane. This mechanism has been observed in the eukaryotic MACPF, perforin-2, where membrane tethering requires structural reorientation to deliver pore-forming activity across distinct membrane compartments [36, 41]. These similarities highlight a conserved strategy among pore forming proteins where localisation and activation are tightly coupled to control toxicity.

In addition to receptor specificity, our findings point to an additional layer of regulation involving proteolytic processing. We discover that bactericidal activity of BSAP-1 depends on proteolytic activation of its N-terminal domain by a surface protease on susceptible cells. This dual-regulatory mechanism, which links target recognition with localized activation, echoes similar strategies seen in pore-forming proteins of both bacterial commensals of the mosquito midgut and the human immune system. While CDCs do not require cleavage for activation, *Elizabethkingia anophelis* and various gut Bacteroidales strains produce CDC-like toxins whose activation depends on proteolytic processing [17]. Similarly, perforin-2, the evolutionary ancestor of eukaryotic MACPF proteins is activated by proteases within the phagophore to restrict toxicity of its pore forming activity [42]. Our data show that together, auxiliary N- and C-terminal domains of BSAP-1 play a key role in controlling activity of the MACPF domain. This functional and evolutionary modularity highlights how BSAP-1, like other soluble toxins, control release of pore-forming residues to prevent off-target activation.

These findings also widen the biological significance of fragipain-family proteases in the gut Bacteroidales. Fragipain was initially characterized as a surface-associated C11 cysteine protease required for maturation of the enterotoxin BFT [25]. Subsequent work showed that fragipain contributes more broadly to shaping the extracellular proteome [26] and can cleave human cell-adhesion molecules on intestinal epithelial cells [26]. Our data now indicate that a fragipain-family protease can also function on susceptible cells to activate BSAP-1. Fragipain has been shown to be necessary for the activity of a distinct *B. fragilis* antibacterial toxin, Fab1, but cleavage was required in the producing cell [16] for its release from the cell. Cleavage of antibacterial toxins on target cells suggests a broader role for surface C11 proteases in Bacteroidales cell biology. Rather than acting only in secretion or maturation of producer-derived effectors, these proteases may also define competitive susceptibility across strains by controlling where and when antibacterial toxins become active.

Our results show how BSAP-1 co-opts proteins on susceptible cells for inter-strain competition; however, the broader role of the bacterial cell envelope remains an open question. Our molecular dynamics simulations show that the BSAP-1 C-terminal domain remains stably associated with OMP-S in a lipid environment, but that the MACPF domain retains considerable orientational flexibility relative to the receptor. This observation is consistent with previous cryo-EM analysis of the detergent-solubilized BSAP-1–OMP-S complex [28], and suggests that receptor binding alone does not orient the pore-forming domain for insertion. One possibility is that receptor binding concentrates BSAP-1 on the target surface, whereas proteolytic cleavage enables a productive orientation or relieves an inhibitory interaction that otherwise prevents oligomerization. Another possibility is that additional features of the native *B. fragilis* outer membrane, including lipid composition or lipopolysaccharide architecture contribute to BSAP-1 activation and assembly. Distinguishing among these models will require future structural analysis in more native membrane contexts.

In summary, our work defines BSAP-1 as a receptor-targeted, protease-activated antibacterial MACPF toxin. The BSAP-1 C-terminal domain mediates selective recognition of OMP-S on susceptible strains where the toxin is locally activated by the proteolytic cleavage of its N-terminal domain. This activation event triggers a BSAP-1 oligomerization and pore formation, both of which are essential for its bactericidal activity. These findings provide a mechanistic framework for understanding how BSAPs combine conserved pore-forming function with target specificity, establishing a general principle by which gut bacteria can spatially restrict toxin activation to shape strain-level competition.

## Acknowledgements

This project has received funding from the British Biological Research Council (UKRI3575) to DB. LEC is supported by the Duchossois Family Institute and R01AI093771 and R01AI181297 from the NIH/NIAID. Research in the SLR group is supported by UKRI Future Leaders Fellowship (MR/Y01975X/1) and Wellcome Trust Discovery Award (301619/Z/23/Z). Cryo-EM data processing used a BBSRC-funded GPU server (BB/X019284/1). We thank Diamond for access and support of cryo-EM facilities accessed through proposal BI39228. We thank S. Islam at Imperial for technical support. All authors declare no financial conflicts of interest.

## Author contributions

SM, DB and LEC, conceptualization; SM (cryoEM), GH (molecular dynamics, formal analysis; SM (cryoEM), GH (molecular dynamics), KF, LEC (bacterial genetics and overlay assays), HN (peptide array and far-western analysis) investigation; SM, DB, GH, LEC and SR data analysis; DB, SR, and LEC, funding acquisition; DB and LC project administration; DB, SR, LEC, supervision; SM, DB validation; SM, GH, visualization; DB and LC writing- original draft; all authors, writing-review and editing.

## Materials and methods

### Bacterial growth and spot agar overlay assay

Strains used and created in this study are listed in Supplementary Table 4. *B. fragilis* strains were grown in supplemented basal medium [43] or on BHIS plates [44] at 37°C under anaerobic conditions. Spot agar overlay assays were performed essentially as described [17]. *B. fragilis* 638RΔ*bfubb* (ΔBF638R_3923) served as the wild type strain and background strain for all mutations. For overlay assays, *B. fragilis* 638RΔ*bfubb* strains were resuspended into supplemented basal medium from a fresh plates to an OD_600_ of 0.25 – 0.3. 5 µl of this suspension was dotted to a BHIS plate and allowed to grow for 20 hours at which time the bacteria were removed with a swab and the plate exposed to chloroform vapor to kill residual bacteria. *B. fragilis* 12905 or *B. fragilis* CL03T12C07 WT or CL03T12C07Δ*fpn* were resuspended from a fresh plate into basal medium to an OD_600_ of ∼0.5 and 100 ul was added to 4 ml top agar and poured over the plate. Zones of growth inhibition were photographed the next day. Overlays using purified His-tagged BSAP-1 were dotted to the plates with 5 µl containing 0.74 µg toxin after the sensitive strains were added in the top agar.

### B. fragilis mutant construction

Site specific mutants in *bsap-1* (BF638R_1646) in *B. fragilis* 638RΔ*bfubb* were made using DNA regions ordered from Genscript that included the altered bases with flanking regions for double cross over recombination. These DNA regions were cloned into pMLS36 [45] and transformed into *E. coli* S17 λ pir. Plasmids were mated into 638RΔ*bfubb* [14] and cointegrates were selected on BHIS plates with 200 µg/ml gentamycin and 10 µg/ml cefoxitin. Cointegrates were passaged in non-selective medium and then plated on BHIS plates with 75 ng/ml anhydrotetracycline. Double recombinants were PCR screened for the mutant genotype and whole genome sequencing was performed to confirm the mutations.

The fragipain gene of *B. fragilis* 638R (Bf638R_2765) and CL03T12C07 (HMPREF1067_01433) were deleted using the primers listed in Supplementary Table 5 and the flanking regions were cloned into BamHI digested pLGB36 [46]. The cointegrate and double crossover methods were performed as listed above except the cointegrates were selected on 200 µg/ml gentamycin and 10 µg/ml erythromycin.

### AlphaFold modelling

AlphaFold 3 (https://alphafoldserver.com/)[18] was used to model BSAP-1 soluble monomer, OMP-S, and OMP-R. AF multimer was used to model the BSAP-1-OMP-S and BSAP-1-OMP-R binding interface. From each run, the top-ranked model (model 0) and corresponding JSON file was mined for confidence values. The per residue predicted aligned error (PAE) was fetched for any residue pair within 6 Å Cα distance, and 8 Å Cα for potential salt bridges. PAE values <5 Å are considered strong confidence, values 5-10 Å are moderate confidence, and those >10 Å are low confidence. For the BSAP-1 chain, the predicted local distance difference test (pLDDT) score was fetched for the terminal 34 residues for BSAP-1 alone or when modelled bound to both OMP-S and OMP-R, with a score closer to 100 increasing confidence.

### Protein expression and purification

His-tagged BSAP-1 constructs were ordered from Genscript so that the N-terminal signal sequence was removed and remainer of the gene or mutant versions were cloned into the NdeI site of pET16b for creation of N-terminally His-tagged proteins. Plasmids were transformed into BL21(DE3) cells and grown in LB medium at 37 °C with orbital shaking at 200 rpm to an OD_600_ of 0.6. BSAP-1 was induced with 0.5 mM isopropyl β-d-1-thiogalactopyranoside (IPTG) and expressed overnight (16 hr). BSAP-1 cell pellets were resuspended in BSAP-1 buffer (Tris 20 mM pH 7.5, 100 mM NaCl) supplemented with cOmplete EDTA-free protease tablets and sonicated on ice. Debris was removed by centrifugation and lysate cleared with a 0.4 μm filter before incubation with Talon Cobalt resin for 1 hr. Resin was washed with BSAP-1 Buffer (10 x column volume), followed by BSAP-1 Buffer + 10 mM imidazole (2 x column volume) before sample was eluted with 200 mM imidazole (8 x column volume). Pooled BSAP-1 fractions were further purified on a Superdex 200 Increase 10/300 GL column (Cytiva) using an AKTA pure system (Cytiva) for structural studies. Sample purity was assessed by SDS-PAGE, pooled, and stored at -70 °C at 1 mg/mL.

Constructs for OMP-S, with its N-terminal signal sequence removed, were ordered from Genscript and cloned into the pET-52b vector with a N-terminal StrepII-Tag tag and a C-terminal His-tag. OMP-S expression was induced with 1 mM IPTG for 3 hr at 37 °C to drive localisation to inclusion bodies. OMP-S cell pellets were resuspended in OMP Resuspension Buffer (Hepes 20 mM pH 8.0, 150 mM NaCl) supplemented with cOmplete EDTA-free protease tablets and sonicated on ice. Inclusion bodies were harvested by first centrifuging (10,000xg) and discarding cell lysate. The pellet was then homogenized with a Dounce Homogenizer in 10 mM Tris pH 8.0, 1 mM EDTA, 1% Triton X-100 and centrifuged (10,000xg). The resulting inclusion body pellet was then washed by homogenizing in 10 mM Tris pH 8.0 before centrifuging again (10,000xg) and harvesting the pellet. To solubilize the inclusion bodies, unfolding buffer (50 mM Tris pH 8.0, 8 M urea) was added and incubated on a shaker overnight (>16 hr). Insoluble material was removed by centrifugation (18,500xg) and the resulting supernatant containing unfolded proteins was incubated with Talon Cobalt resin for 1 hr. Resin was then washed with unfolding buffer (20 x column volume), unfolding buffer + 10 mM imidazole (2 x column volume) before being eluted with unfolding buffer + 200 mM imidazole (10 x column volume). Elutions containing OMP-S were concentrated before being added dropwise to 20-fold v/v refolding buffer (200 mM CAPS, 400 mM NaCl, 50 mM Tris, 0.5% LDAO, pH 11) and left spinning overnight. The pH was adjusted to 8.0 and clarified with a 0.4 μm filter, before being concentrated and purified with a Superdex 200 Increase 10/300 GL column (Cytiva) pre-equilibrated with 50 mM Tris pH 7.5, 150 mM NaCl with either 0.1% LDAO or 0.05% DDM depending on downstream application.

### Peptide array: far-western blot

Peptide arrays were synthesised on derivatised cellulose membranes using a ResPepSL Peptide Synthesiser (CEM). The membranes contained 8–10 ethylene glycol spacer units between the cellulose and the terminal amino group, with a loading of approximately 400 nmol/cm². Peptides were assembled by standard Fmoc solid-phase synthesis using DIC/Oxyma-mediated coupling. Fmoc groups were removed with 20% piperidine in DMF containing 1% formic acid between coupling cycles. After synthesis, side-chain protecting groups were removed by treatment with 95% trifluoroacetic acid, 3% triisopropylsilane and 2% water for 4 h. Membranes were then washed sequentially with dichloromethane, ethanol and water, air-dried and stored at −20°C until use. For far-western blotting, membranes were first blocked with 5% milk TBS-T before incubation with 0.2 μM OMP-S in 0.1% LDAO for 2.5 hours. Unbound protein was washed from the membrane and incubated with 1:2000 6x-His Tag Monoclonal Antibody (MA1-21315, Thermofisher). Bound protein was detected using 1:10000 HRP-conjugated goat anti-mouse IgG secondary antibody (31430, Thermofisher) and visualised by enhanced chemiluminescence. Values for each spot were normalised to WT peptide for each lane in Fiji [47].

### BSAP-1 pore assembly on liposomes

1,2-dioleoyl-sn-glycero-3-phosphocholine (DOPC) lipid films were rehydrated to 5 mg/ml in BSAP-1 Buffer (Tris 20 mM pH 7.5, 100 mM NaCl) and subjected to 4 x freeze thaw cycles, before being extruded through a 100 nm polycarbonate membrane with an Avanti Mini Extruder. Liposomes were then diluted to 1 mg/mL and incubated with BSAP-1 100 µg/mL at room temperature for 15 minutes. BSAP-1 was activated with 2 µg/mL Trypsin (from bovine pancreas, Sigma) and incubated at 37 °C, 15 minutes, before the reaction was stopped with 10 µg/mL Soybean Trypsin Inhibitor. Samples were then centrifuged (60,000xg) to pellet liposomes and remove soluble material. Liposomes were resuspended at 0.5 mg/mL and the presence of BSAP-1 fragments and oligomers were assessed using SDS-PAGE.

### Cryo-EM grid preparation and data collection

Lacey carbon Au 300-mesh grids (Electron Microscopy Sciences) were glow discharged for 45 seconds using a Cressington 208 (Cressington Scientific Instruments). 4 µL of liposome assembled BSAP-1 pores was applied to the carbon film and incubated for 60 seconds and blotted (first application). 4 µL sample was then reapplied, blotted, and plunge frozen into liquid ethane using a Vitrobot mark IV (Thermo fisher Scientific) with a blot time of 6.5 seconds and a blot force of -2. Data were collected at the electron Bio-Imaging centre (eBIC) using a Titan Krios (Thermo Fisher Scientific) operating at 300 kV equipped with a Falcon 4i direct electron detector with a Selectris X energy filter (20 eV). Grids were imaged at a nominal magnification of 130,000x (0.931 Å/pixel).

### Cryo-EM image processing

8,780 movies were imported into CryoSPARC v4.7.0 [48] and corrected for beam induced motion using patch motion correction, and CTF parameters estimated using patch CTF. Particles were manually picked from a random sub-selection of micrographs at a range of defocus and used to custom train models in Topaz v2.0.5 [49]. Particles were extracted with a box size of 336 px, downsampled by 4, and cleaned using 2D classification. Multiple rounds of multi-class ab initio reconstructions were then performed to further remove junk particles. Particles from the best ab initio classes underwent homogenous refinements. Duplicate particles were then removed and re-extracted without downsampling and individually corrected using local CTF refinement. A mask covering the pore and excluding the lipid membrane was used for local refinement in C1 revealing 13 monomers which was further refined using C13 symmetry.

### Model building and refinement

An initial model of the BSAP-1 pore was obtained by running Modelangelo implemented in Relion 5 [50], with the sharpened C13 map and BSAP-1 FASTA sequence (minus the N-terminus) as inputs. Beta hairpins that were truncated by Modelangelo were rebuilt using Modeller v.10.7 [51]. The best monomer model was then refined into the EM density using Isolde v1.9 [52]. Atomic model refinement was then performed using Servalcat with the REFMAC5 engine [53], operated through the Doppio pipeline within the CCPEM software suite [54]. A 13-mer was generated using symmetry expansion with restrained symmetry to best fit the data using half maps. Map-model FSC and full cryo-EM validation were assessed using inbuilt validation tools (Supplementary Table 3). The pore form and soluble forms of BSAP-1 were visualised in ChimeraX (v.1.9) [55] and compared using the matchmaker command.

### BSAP pore homology model creation and membrane visualisation

BSAP-2, BSAP-3, and BSAP-4 monomers in the soluble state were predicted using the AF3 server. Monomers in the pore state were predicted using colabfold [56] with the BSAP-1 pore monomer pdb as a template. Correct domain boundaries were confirmed by comparison with the soluble domains and clustalW multiple sequence alignment [57]. Secondary structures of initial models were shifted into place using the BSAP-1 unsharpened cryoEM map with Isolde. Full 13-mer pores for each BSAP were generated using symmetry expansion in Servalcat with the BSAP-1 map low-pass filtered to 8 Å.

### Atomistic molecular dynamics simulations

All simulations were performed using GROMACS v2023.3 and v2024.3 [58]. The AF3 model of BSAP-1 bound to OMP-S was used as the starting coordinates, with OMP-S lacking the signal sequence (residues 28-560) and BSAP-1 with the N-terminus removed (residues 37-372). The atomistic model was energy minimised, and embedded into an 18 nm by 18 nm dipalmitoylphosphatidylcholine (DOPC) lipid bilayer using the standard CHARMM-GUI membrane builder protocol [59]. The system was solvated with TIP3P waters and neutralised to a concentration of 0.15 M NaCl. Further rounds of energy minimisation and equilibration were carried out in which position restraints were faded out in a stepwise manner over the standard 6-step CHARMM-GUI protocol for GROMACS simulations [60]. Atomistic simulations were performed using the CHARMM36m [61] and CHARMM36 [62] force fields. The LINCS algorithm was used to constrain covalent bond lengths [63]. Long-range electrostatics were described using Particle Mesh Ewald (PME) [64]. Van der Waals interactions were calculated using the Verlet method with a 12 Å cutoff. Equations of motion were integrated using a 2 fs time step. Periodic boundary conditions were applied. Temperature was controlled at 303.15 K using a V-rescale thermostat [65] (*τ_t_* = 1.0). Pressure was maintained at 1 bar using a C-rescale barostat [66] (*τ_p_* = 5.0) with semi-isotropic coupling and a compressibility of 4.5 x 10^-5^ bar^-1^ [58]. Simulations were analysed using tools available in GROMACS, Visual Molecular Dynamics (VMD) [67], PyMOL [68], and PyLipID [69].

## Data availability

Data supporting the findings of this manuscript are available from the corresponding authors upon reasonable request. BSAP-1 pore cryo-EM map and corresponding structural model is available in the Electron microscopy database EMD-58950 and Protein Data Bank PDB: 32KD, respectively. Molecular dynamics simulation data files used in this study are uploaded to Zenodo database and will be released on publication.

## References

1. Faith, J.J., et al., The long-term stability of the human gut microbiota. Science, 2013. 341(6141): p. 1237439.

2. Wexler, H.M., Bacteroides: the good, the bad, and the nitty-gritty. Clin Microbiol Rev, 2007. 20(4): p. 593–621.

3. Van Tassell, R.L., D.M. Lyerly, and T.D. Wilkins, Purification and characterization of an enterotoxin from Bacteroides fragilis. Infect Immun, 1992. 60(4): p. 1343–50.

4. San Joaquin, V.H., et al., Association of Bacteroides fragilis with childhood diarrhea. Scand J Infect Dis, 1995. 27(3): p. 211–5.

5. Rhee, K.J., et al., Induction of persistent colitis by a human commensal, enterotoxigenic Bacteroides fragilis, in wild-type C57BL/6 mice. Infect Immun, 2009. 77(4): p. 1708–18.

6. Chung, L., et al., Bacteroides fragilis Toxin Coordinates a Pro-carcinogenic Inflammatory Cascade via Targeting of Colonic Epithelial Cells. Cell Host Microbe, 2018. 23(2): p. 203–214 e5.

7. Valguarnera, E. and J.B. Wardenburg, Good Gone Bad: One Toxin Away From Disease for Bacteroides fragilis. J Mol Biol, 2020. 432(4): p. 765–785.

8. Sun, L., et al., Gut microbiota and intestinal FXR mediate the clinical benefits of metformin. Nat Med, 2018. 24(12): p. 1919–1929.

9. Xia, Y., et al., Bacteroides Fragilis in the gut microbiomes of Alzheimer’s disease activates microglia and triggers pathogenesis in neuronal C/EBPbeta transgenic mice. Nat Commun, 2023. 14(1): p. 5471.

10. Coyne, M.J. and L.E. Comstock, Type VI Secretion Systems and the Gut Microbiota. Microbiol Spectr, 2019. 7(2).

11. Chatzidaki-Livanis, M., M.J. Coyne, and L.E. Comstock, An antimicrobial protein of the gut symbiont Bacteroides fragilis with a MACPF domain of host immune proteins. Mol Microbiol, 2014. 94(6): p. 1361–74.

12. McEneany, V.L., et al., Acquisition of MACPF domain-encoding genes is the main contributor to LPS glycan diversity in gut Bacteroides species. ISME J, 2018. 12(12): p. 2919–2928.

13. Shumaker, A.M., et al., Identification of a Fifth Antibacterial Toxin Produced by a Single Bacteroides fragilis Strain. J Bacteriol, 2019. 201(8).

14. Chatzidaki-Livanis, M., et al., Gut Symbiont Bacteroides fragilis Secretes a Eukaryotic-Like Ubiquitin Protein That Mediates Intraspecies Antagonism. mBio, 2017. 8(6).

15. Coyne, M.J., et al., A family of anti-Bacteroidales peptide toxins wide-spread in the human gut microbiota. Nat Commun, 2019. 10(1): p. 3460.

16. Bao, Y., et al., A Common Pathway for Activation of Host-Targeting and Bacteria-Targeting Toxins in Human Intestinal Bacteria. mBio, 2021. 12(4): p. e0065621.

17. Evans, J.C., et al., A proteolytically activated antimicrobial toxin encoded on a mobile plasmid of Bacteroidales induces a protective response. Nat Commun, 2022. 13(1): p. 4258.

18. Abrahamsen, H.L., et al., Distant relatives of a eukaryotic cell-specific toxin family evolved a complement-like mechanism to kill bacteria. Nat Commun, 2024. 15(1): p. 5028.

19. Stevens, L.M., et al., Localized requirement for torso-like expression in follicle cells for development of terminal anlagen of the Drosophila embryo. Nature, 1990. 346(6285): p. 660–3.

20. Ni, T., K. Harlos, and R. Gilbert, Structure of astrotactin-2: a conserved vertebrate-specific and perforin-like membrane protein involved in neuronal development. Open Biol, 2016. 6(5).

21. Ivanova, M.E., et al., The pore conformation of lymphocyte perforin. Sci Adv, 2022. 8(6): p. eabk3147.

22. Serna, M., et al., Structural basis of complement membrane attack complex formation. Nat Commun, 2016. 7: p. 10587.

23. Yu, X., et al., Cryo-EM structures of perforin-2 in isolation and assembled on a membrane suggest a mechanism for pore formation. EMBO J, 2022. 41(23): p. e111857.

24. Roelofs, K.G., et al., Bacteroidales Secreted Antimicrobial Proteins Target Surface Molecules Necessary for Gut Colonization and Mediate Competition In Vivo. mBio, 2016. 7(4).

25. Choi, V.M., et al., Activation of Bacteroides fragilis toxin by a novel bacterial protease contributes to anaerobic sepsis in mice. Nat Med, 2016. 22(5): p. 563–7.

26. Pierce, J.V., et al., A clostripain-like protease plays a major role in generating the secretome of enterotoxigenic Bacteroides fragilis. Mol Microbiol, 2021. 115(2): p. 290–304.

27. Herrou, J., et al., Activation Mechanism of the Bacteroides fragilis Cysteine Peptidase, Fragipain. Biochemistry, 2016. 55(29): p. 4077–84.

28. Borgini, S., et al., Identification of receptor-binding domains of Bacteroidales antibacterial pore-forming toxins. J Biol Chem, 2026. 302(2): p. 111113.

29. Abramson, J., et al., Accurate structure prediction of biomolecular interactions with AlphaFold 3. Nature, 2024. 630(8016): p. 493–500.

30. Hotze, E.M., et al., Monomer-monomer interactions drive the prepore to pore conversion of a beta-barrel-forming cholesterol-dependent cytolysin. J Biol Chem, 2002. 277(13): p. 11597–605.

31. Ramachandran, R., R.K. Tweten, and A.E. Johnson, Membrane-dependent conformational changes initiate cholesterol-dependent cytolysin oligomerization and intersubunit beta-strand alignment. Nat Struct Mol Biol, 2004. 11(8): p. 697–705.

32. Johnstone, B.A., et al., Structural basis for the pore-forming activity of a complement-like toxin. Sci Adv, 2025. 11(13): p. eadt2127.

33. Menny, A., et al., CryoEM reveals how the complement membrane attack complex ruptures lipid bilayers. Nat Commun, 2018. 9(1): p. 5316.

34. van Pee, K., et al., CryoEM structures of membrane pore and prepore complex reveal cytolytic mechanism of Pneumolysin. Elife, 2017. 6.

35. Shatursky, O., et al., The mechanism of membrane insertion for a cholesterol-dependent cytolysin: a novel paradigm for pore-forming toxins. Cell, 1999. 99(3): p. 293–9.

36. Ni, T., et al., Structure and mechanism of bactericidal mammalian perforin-2, an ancient agent of innate immunity. Sci Adv, 2020. 6(5): p. eaax8286.

37. Elhenawy, W., M.O. Debelyy, and M.F. Feldman, Preferential packing of acidic glycosidases and proteases into Bacteroides outer membrane vesicles. mBio, 2014. 5(2): p. e00909–14.

38. Rakoff-Nahoum, S., M.J. Coyne, and L.E. Comstock, An ecological network of polysaccharide utilization among human intestinal symbionts. Curr Biol, 2014. 24(1): p. 40–49.

39. Shen, Y., et al., Outer membrane vesicles of a human commensal mediate immune regulation and disease protection. Cell Host Microbe, 2012. 12(4): p. 509–20.

40. Sheikh, A., et al., Outer Membrane Vesicles From Bacteroides fragilis Contain Coding and Non-Coding Small RNA Species That Modulate Inflammatory Signalling in Intestinal Epithelial Cells. J Extracell Biol, 2025. 4(10): p. e70086.

41. Jiao, F., et al., Perforin-2 clockwise hand-over-hand pre-pore to pore transition mechanism. Nat Commun, 2022. 13(1): p. 5039.

42. Rodriguez-Silvestre, P., et al., Perforin-2 is a pore-forming effector of endocytic escape in cross-presenting dendritic cells. Science, 2023. 380(6651): p. 1258–1265.

43. Pantosti, A., et al., Immunochemical characterization of two surface polysaccharides of Bacteroides fragilis. Infect Immun, 1991. 59(6): p. 2075–82.

44. Garcia-Bayona, L. and L.E. Comstock, Streamlined Genetic Manipulation of Diverse Bacteroides and Parabacteroides Isolates from the Human Gut Microbiota. mBio, 2019. 10(4).

45. Sheahan, M.L., et al., A ubiquitous mobile genetic element changes the antagonistic weaponry of a human gut symbiont. Science, 2024. 386(6720): p. 414–420.

46. Ito, T., et al., Genetic and Biochemical Analysis of Anaerobic Respiration in Bacteroides fragilis and Its Importance In Vivo. mBio, 2020. 11(1).

47. Schindelin, J., et al., Fiji: an open-source platform for biological-image analysis. Nat Methods, 2012. 9(7): p. 676–82.

48. Punjani, A., et al., cryoSPARC: algorithms for rapid unsupervised cryo-EM structure determination. Nat Methods, 2017. 14(3): p. 290–296.

49. Bepler, T., et al., Positive-unlabeled convolutional neural networks for particle picking in cryo-electron micrographs. Nat Methods, 2019. 16(11): p. 1153–1160.

50. Jamali, K., et al., Automated model building and protein identification in cryo-EM maps. Nature, 2024. 628(8007): p. 450–457.

51. Eswar, N., et al., Comparative protein structure modeling using Modeller. Curr Protoc Bioinformatics, 2006. Chapter 5: p. Unit-5 6.

52. Croll, T.I., ISOLDE: a physically realistic environment for model building into low-resolution electron-density maps. Acta Crystallogr D Struct Biol, 2018. 74(Pt 6): p. 519–530.

53. Yamashita, K., et al., Cryo-EM single-particle structure refinement and map calculation using Servalcat. Acta Crystallogr D Struct Biol, 2021. 77(Pt 10): p. 1282–1291.

54. Burnley, T., C.M. Palmer, and M. Winn, Recent developments in the CCP-EM software suite. Acta Crystallogr D Struct Biol, 2017. 73(Pt 6): p. 469–477.

55. Pettersen, E.F., et al., UCSF ChimeraX: Structure visualization for researchers, educators, and developers. Protein Sci, 2021. 30(1): p. 70–82.

56. Mirdita, M., et al., ColabFold: making protein folding accessible to all. Nat Methods, 2022. 19(6): p. 679–682.

57. Thompson, J.D., T.J. Gibson, and D.G. Higgins, Multiple sequence alignment using ClustalW and ClustalX. Curr Protoc Bioinformatics, 2002. Chapter 2: p. Unit 2 3.

58. Abraham, M.J.M., T.; Schulz, R.; Páll, S.; Smith, J.C.; Hess, B.; Lindahl, E., GROMACS: High performance molecular simulations through multi-level parallelism from laptops to supercomputers. SoftwareX, 2015. 1–2: p. Pages 19-25.

59. Wu, E.L., et al., CHARMM-GUI Membrane Builder toward realistic biological membrane simulations. J Comput Chem, 2014. 35(27): p. 1997–2004.

60. Lee, J., et al., CHARMM-GUI Input Generator for NAMD, GROMACS, AMBER, OpenMM, and CHARMM/OpenMM Simulations Using the CHARMM36 Additive Force Field. J Chem Theory Comput, 2016. 12(1): p. 405–13.

61. Huang, J., et al., CHARMM36m: an improved force field for folded and intrinsically disordered proteins. Nat Methods, 2017. 14(1): p. 71–73.

62. Kako, K.J. and S.C. Vasdev, Effects of a high fat--high erucic acid diet on the lipid metabolism and contractility of the rat heart. Biochem Med, 1979. 22(1): p. 76–87.

63. Hess, B., et al., LINCS: A linear constraint solver for molecular simulations. Journal of Computational Chemistry, 1997. 18(12): p. 1463–1472.

64. Essmann, U., et al., A smooth particle mesh Ewald method. The Journal of Chemical Physics, 1995. 103(19): p. 8577–8593.

65. Bussi, G., D. Donadio, and M. Parrinello, Canonical sampling through velocity rescaling. The Journal of Chemical Physics, 2007. 126(1).

66. Bernetti, M. and G. Bussi, Pressure control using stochastic cell rescaling. The Journal of Chemical Physics, 2020. 153(11).

67. Humphrey, W.D., A.; Schulten, K., VMD: Visual molecular dynamics. Journal of Molecular Graphics, 1996. 14(1): p. 33–38.

68. Schrodinger, L.L.C., The PyMOL Molecular Graphics System. 2024.

69. Song, W., et al., PyLipID: A Python Package for Analysis of Protein-Lipid Interactions from Molecular Dynamics Simulations. J Chem Theory Comput, 2022. 18(2): p. 1188–1201.

