## Supplementary information for "Bacteroidales Secreted Antimicrobial Protein-1 receptor recognition is coupled to protease-activation to form bactericidal pores"

Shannon N. Mostyn[1], Katia Flores[2], George Hedger[1], Hema Nagaraj[3], Sarah L. Rouse[1], Laurie E. Comstock[2*], Doryen Bubeck[1*]

[1]Department of Life Sciences, Imperial College, London, UK

[2]Duchossois Family Institute, Department of Microbiology, University of Chicago, Chicago, Illinois, USA

[3]Chemical Biology STP, The Francis Crick Institute, London, UK

**Supplementary information**

Supplementary Figures 1-7

Supplementary Tables 1-5

Supplementary References 1-5

### Supplementary Figures

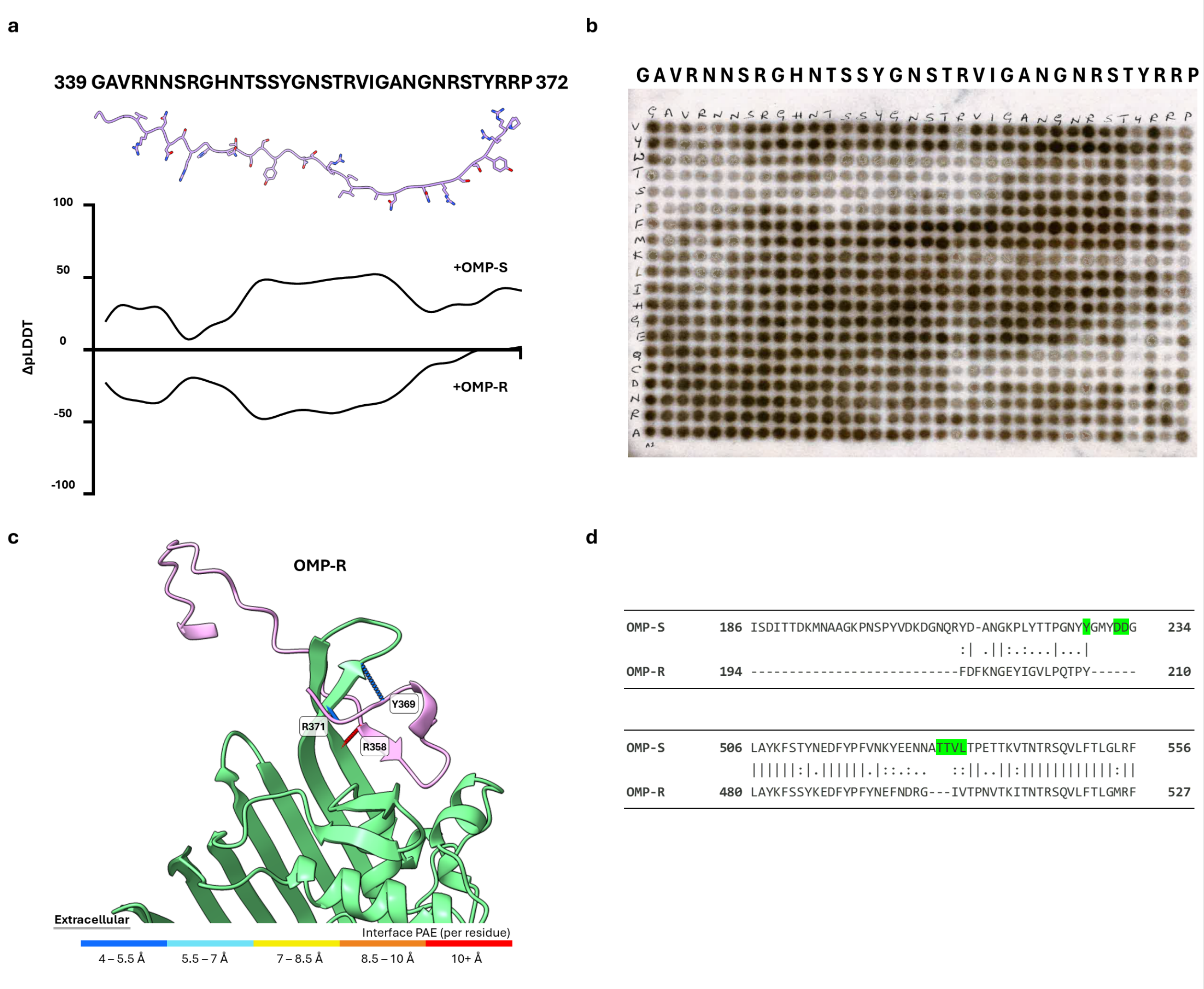

**Supplementary Figure 1. a)** Per-residue pLDDT of predicted BSAP-1 C-terminal domain (residues 339-372) interactions with OMP-S and OMP-R. pLDDT scores were normalised to values for the AF3 BSAP-1 monomer prediction (ΔpLDDT). **b)** Raw peptide array. Peptides corresponding to the BSAP-1 C-terminal domain (residues 339-372) with all possible amino acid substitutions were constructed on a peptide array and probed for His-tagged OMP-S binding via far-Western blotting. **c)** AF3 predicted binding interface of OMP-R and BSAP-1. PAE scores of the only 3 residues which are within 5 Å (Supplementary Table 1b) are indicated. **d)** Sequence alignment of OMP-S and OMP-R showing differences in the extracellular loops. Specific interface residues identified from the OMP-S-BSAP-1 modelling highlighted green.

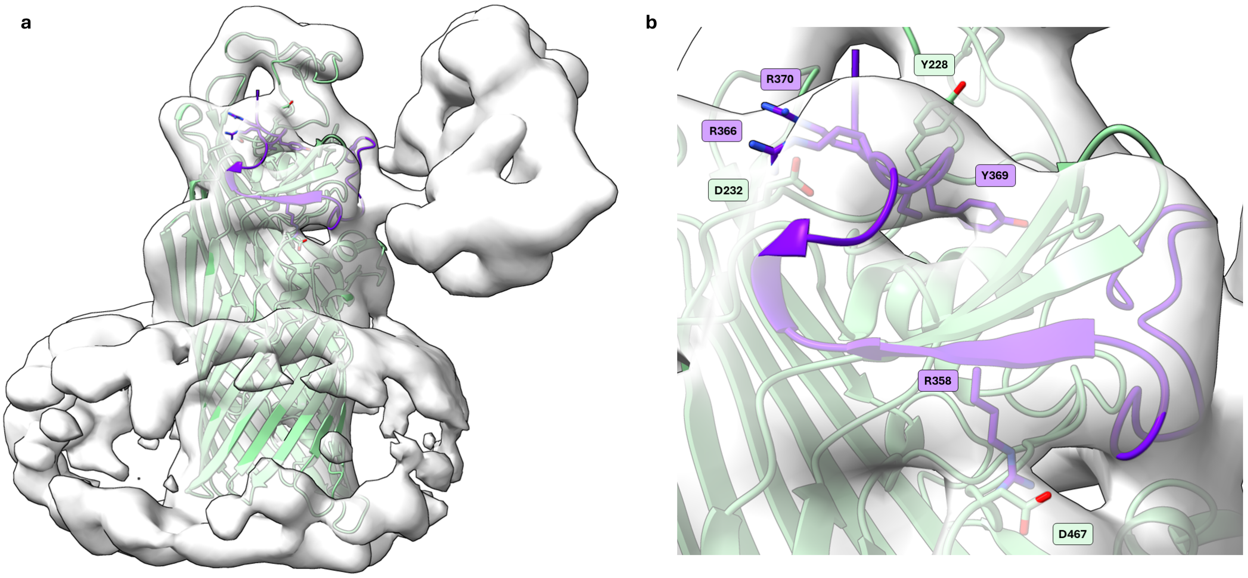

**Supplementary Figure 2. a)** AF3 BSAP-1 (purple)-OMP-S (green) structural model docked into EMD-53314 (transparent surface)[1] with the voxel size adjusted to 1.506 Å/pixel. Model for BSAP-1 MACPF is hidden for clarity. **b)** Close up of predicted binding interface. Residues whose sidechains contribute to the interface are shown.

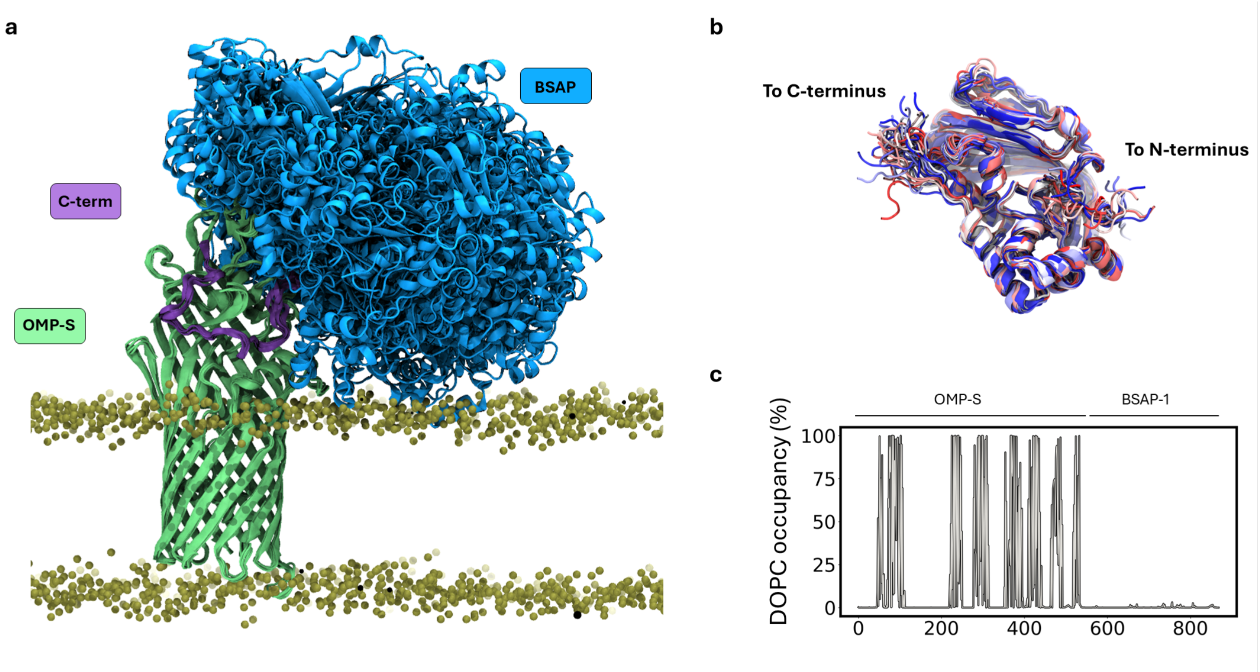

**Supplementary Figure 3. Molecular dynamics simulation of BSAP-1 (ΔN-term) in complex with OMP-S. a)** Overlay of 50 evenly distributed structures over combined 5 x 300 ns ensemble. **b)** Overlay of the BSAP-1 MACPF domain across the simulation shows no major conformational changes when engaged with OMP-S. MACPF domain of BSAP-1 is coloured red-white-blue according to simulation time. N- and C-terminal boundaries of the MACPF domain are indicated. **c)** Lipid occupancy values for individual residues of OMP-S and BSAP-1.

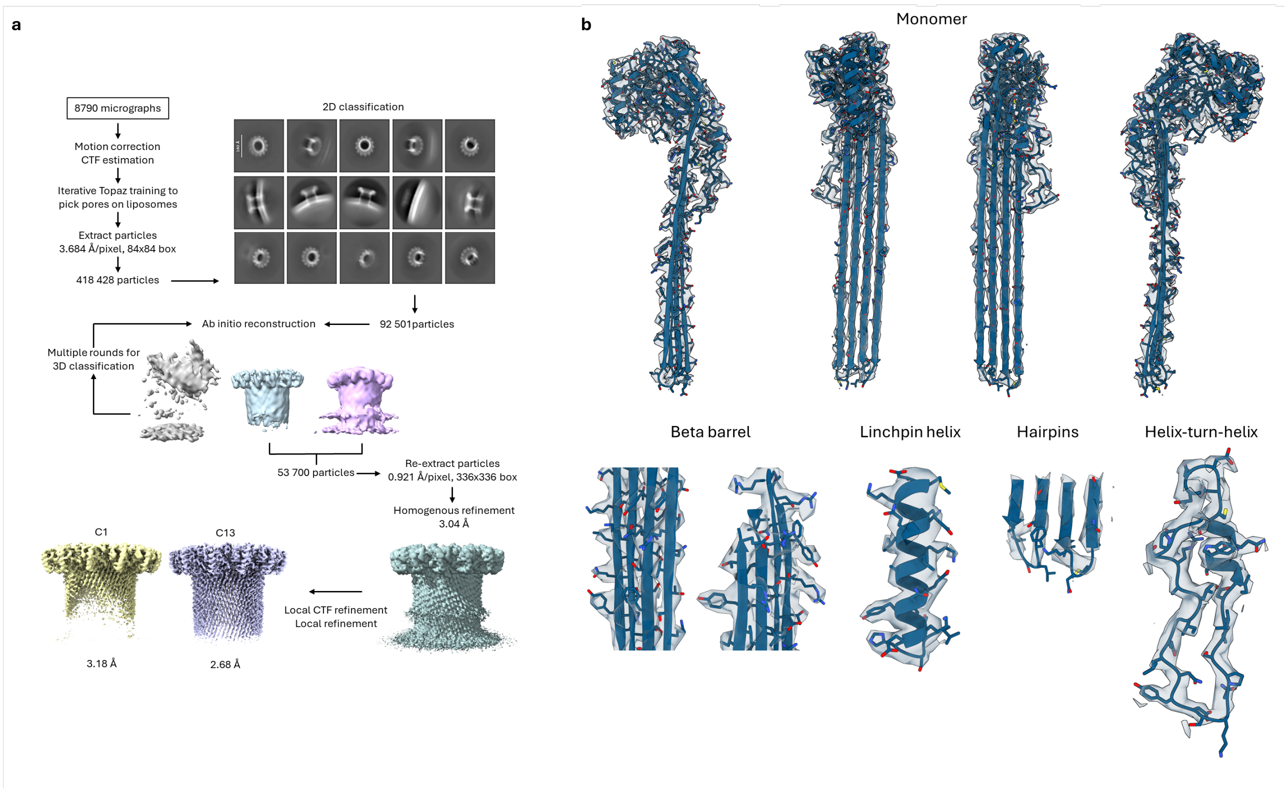

**Supplementary Figure 4. a)** BSAP-1 pore data processing pipeline. **b)** Map-model fit. Threshold to 0.0624 σ (monomer, hairpins), and 0.0992 σ (beta barrel, linchpin helix, helix-turn-helix).

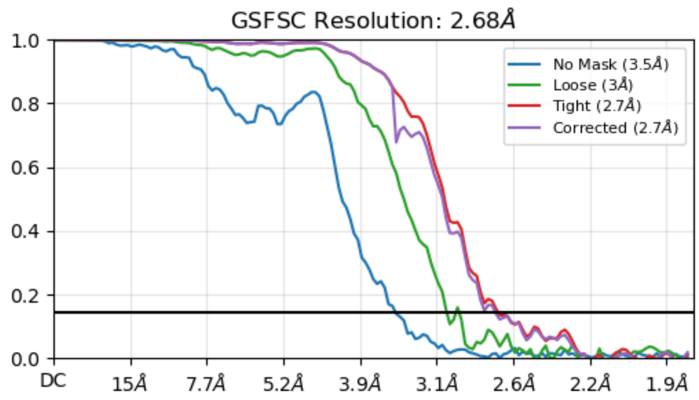

**Supplementary Figure 5.** BSAP-1 pore map FSC (gold standard cutoff 0.143).

```
BSAP-1      --------------MKKLGLLLMLFT-----FAACTNDD-FSVLQENE-------------EITLPTAVTRASGDKLYDLLGYGYDVTGL    58
BSAP-2      --------------MKKKHFLFIPIITMGIFAISCTNTFDEQFVQEDQ-----------------QMPTTKSMGDGQYQVLGYGYDITGE    64
BSAP-3      --------------MRKINLCYFL--FV-GVLVSCTNTYDVCEMQRNE----------------TEDMSSRMAGDGVYDVLGFGYDITED    61
BSAP-4      MKREEISYKNYSKLLIKLSLLGIS-----FIIVSCNKTDLITDNPNNELGQLPQENGITIKTITYSPQITRSSGDGANDLIGYGYNCLYA    85
                          : *  :  :          :*.:        .::                     ::  **   :::*:**:

BSAP-1      Y-FTSASAKSKIIDIVALRKDYEERVDIGAVPSNYARMTSGTTAQDYTRNVTSKVKLGGALSL----------FSG-------------    122
BSAP-2      Y-LARQSIKNEVLDIAGFYAKNPQLYKVEYVGEIKDNFYAGEDYMNYVKDIINKTNFSGSVAAKINPKKD-GATSTDTSNK---EVPYSL    144
BSAP-3      Y-LGENSTRLQVIDVKALIRDNGDRFYNPFIGTIDQKIVAGEDAQSFLQNVITDSNFKGSVAAMDIKKDAEGFFSG-------------    132
BSAP-4      TGDNPNGAANKVIDLKRFESGVGID---PMT---------G-EKKIFPKGIIEESIMHGDVTATEIFEKSKKEFKKSISAKGKLKLGFLP    162
                  .   :::*:  :                      *     : :.:  .  : * ::            .

BSAP-1      --SLSSSF--SSTQHYTSKYSIADYTSFIRRRRLFLTA-STELLSKYLTKMFVDDLSKQSPSFIIQHYGTHVLTDITLGGRITVLYRSSI    207
BSAP-2      SANVSFDKESSNKTSITTKETYIRVDQIKQLERYQLYA-DPTVLADYLTPYFKSRLSTASPDAIIEEFGTHILIDFNVGGRLSIFYKSTI    233
BSAP-3      --TLSAGFKSASKYSYSSKYSFAKAEVTKKQRKYFVDA-DVELLSRYLSSNFVSDLGKYSADQIVAKYGTHVLTNITVGGSYIAYYKSAI    219
BSAP-4      WGGIQFSAEYDSNFETKDEYSFYKMDIIRNVRRLYLSSSSIERLKYFLTDEFRYALKHASGTEIINDYGTHVLVDFVLGGKLSVVTSAID    252
               :. .    ..   . : :        . .:  : : .   *  :*:  *   *   *   *: .:***:* :: :**       :

BSAP-1      NTSKKTA----TVEAGCASGIKNMFNLSVDGHYDQTLVKDNSE----QEIVYRTEGGDPSRALIGQLNYDS-KNPSVIDISSWQQSCDDN    288
BSAP-2      TDNLKVESKTKIAKGG-ITGAIKTVNLSFSGSSTTTEVEQYQRRNSNWSCNVNMYGGQHDGHTI-TITSDG-ATNHTFNLSGWQQSVDKT    320
BSAP-3      IEENTGTEKKQTVSAG-AKYNMSKVGLDFDGTWSTTTITEANKKNSDWTCTIQCVGGTTSGTTI-TLAPNQ-GPTTTINIGAWSQSVDDT    306
BSAP-4      HSDNKDELFKF------STKIFKIISASASSS----ISKSNYLK--NTSLTILQAGGSEIQAVKRYIEEDGTINSDIFNYKEWIKSINKE    330
            . .             .   . .. . .  .        .               **         :  :       ::   * :* :.

BSAP-1      NMTLVDAEPGSLIPIYDLVSDMGKKEQLKL AVENYLKEHS--------YVDIDDSSKGGA    340
BSAP-2      HCVLTEINFNKTYPIYEFIKDPIKKQQIKD AAEKYIKSKI--------RPIIEVKPMFQI    372
BSAP-3      HSRLVDVDWNATYPIYDLVSDPVKKADLKL AVEKYINSKK--------ISVIKLVTLYQL    358
BSAP-4      TSVLVASDTKKMIGIWELTDDPKIKTKILA ELEKRGNNLGIIGSKSGEIVTGKSTPMYKL    390
               *.  :      *::: .*   * .:    *:  :.              .
```

**Supplementary Figure 6. Sequence alignment of BSAP N-terminal and MACPF domains.**

Residues that undergo the helix-to-hairpin transition during pore formation: helical bundle 1 (HB1, cyan) and helical bundle 2 (HB2, green) are colored as defined by AF3 predicted soluble structures. Fragipain cleavage site is highlighted in red. C-terminal domain sequences are omitted.

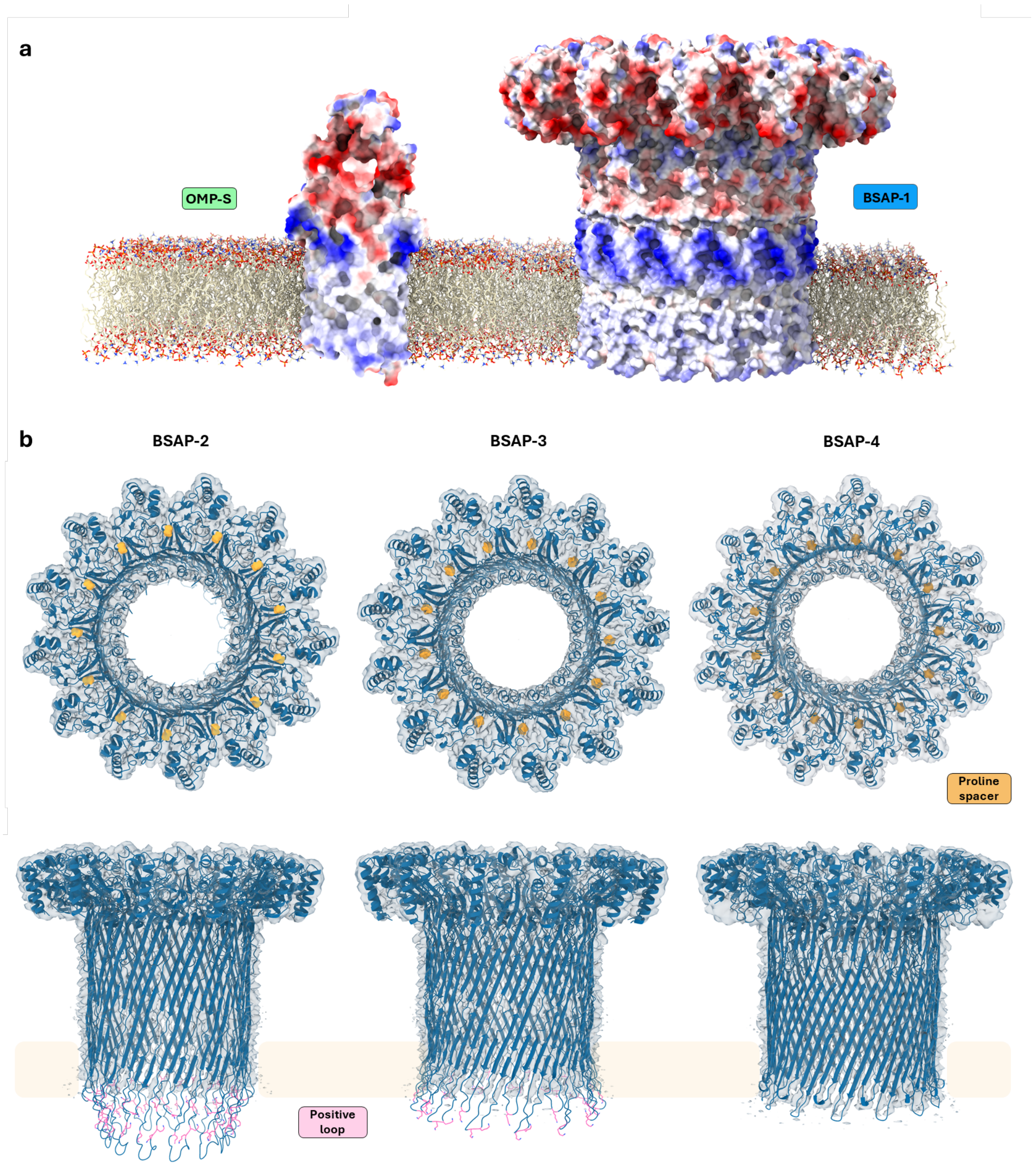

**Supplementary Figure 7. a)** Coulombic electrostatic potential ranging from −10 (red) to 10 (blue) kcal/(mol·e) calculated from the AF3 OMP-S model and the model derived from the cryo-EM structure BSAP-1 pore (this study) embedded in a DOPC bilayer with PPM 3.0 [2]. **b)** AF models of BSAP homologues (BSAP-2, BSAP-3, BSAP-4) using the model for BSAP-1 pore as a template and fitted into the cryoEM structure of BSAP-1 (transparent surface). Structurally conserved proline spacer residue which lies at the oligomerization interface is shown (yellow). Longer charged hairpins of glycan-binding BSAPs (BSAP-2 and BSAP-3) are colored pink.

### Supplementary Tables

**Supplementary Table 1. Analysis of per residues PAE distances for residues <6 Å (backbone) and <8.0 Å (sidechain) (a-b) and pLDDT scores of BSAP-1 C-terminus (c) for AF3 predicted models.**

**a Per residue PAE score for the AF3 OMP-S-BSAP1 model**

| OMP-S | BSAP-1 | Distance (Å) | PAE (Å) |
|---|---|---:|---:|
| G145 | A362 | 5.44 | 8.4 |
| N146 | A362 | 5.83 | 8.2 |
| N146 | R366 | 5.81 | 7.4 |
| Y151 | Y369 | 5.35 | 5.2 |
| G159 | R342 | 5.20 | 5.4 |
| G159 | N343 | 5.03 | 6.5 |
| G159 | S345 | 5.73 | 8.7 |
| I160 | R342 | 4.37 | 6.4 |
| T161 | V341 | 5.81 | 5.7 |
| T161 | R342 | 5.83 | 6.1 |
| K162 | A340 | 5.48 | 6.3 |
| K162 | V341 | 4.92 | 5.3 |
| W163 | A340 | 4.72 | 6.1 |
| G164 | G338 | 4.93 | 8.1 |
| G164 | G339 | 4.72 | 7.6 |
| G164 | A340 | 4.29 | 7.2 |
| D165 | G338 | 5.31 | 7.9 |
| Y228 | Y369 | 4.68 | 4.9 |
| Y228 | R370 | 5.73 | 5.6 |
| Y228 | R371 | 5.03 | 5.2 |
| G229 | Y369 | 5.57 | 5.1 |
| G229 | R370 | 5.53 | 5.6 |
| G229 | R371 | 5.52 | 4.5 |
| M230 | R371 | 5.47 | 4.7 |
| Y231 | R370 | 5.78 | 5 |
| D232 | R366 | 7.80 | 7.8 |
| D232 | R370 | 7.00 | 5.0 |
| F425 | N349 | 5.39 | 6.3 |
| F425 | T350 | 4.72 | 6 |
| S428 | T350 | 5.62 | 5.9 |
| S428 | S351 | 5.82 | 5.9 |
| T429 | S351 | 5.95 | 6.2 |
| D467 | R338 | 6.90 | 5.6 |

| | | | |
|---|---|---|---|
| Y474 | G347 | 5.77 | 9 |
| T531 | S356 | 4.85 | 5.2 |
| T532 | S356 | 5.49 | 6 |
| T532 | T357 | 4.90 | 6 |
| V533 | T357 | 5.97 | 5.7 |
| V533 | R358 | 4.69 | 4.8 |
| L534 | V359 | 4.83 | 5.9 |
| L534 | I360 | 5.89 | 6.3 |
| T535 | I360 | 5.16 | 4.1 |
| P536 | I360 | 5.98 | 4.8 |
| P536 | G361 | 4.73 | 5.4 |
| E537 | I360 | 5.71 | 5.6 |
| E537 | I361 | 4.35 | 6.5 |

**b Per residue PAE score for the AF3 OMP-R-BSAP1 model**

| OMP-R | BSAP-1 | Distance (Å) | PAE (Å) |
|---|---|---|---|
| S249 | H235 | 5.99 | 25.4 |
| M250 | H235 | 5.31 | 25.4 |
| S293 | H235 | 5.55 | 25.6 |
| M302 | Y369 | 5.96 | 4.5 |
| M302 | R370 | 5.80 | 5.5 |
| M302 | R371 | 4.95 | 4.8 |
| M302 | P372 | 5.89 | 6.4 |
| H303 | Y369 | 5.49 | 5.1 |
| H303 | R370 | 5.22 | 5.3 |
| H303 | R371 | 5.71 | 6.1 |
| H303 | P372 | 5.18 | 7.8 |
| N304 | T368 | 5.95 | 6.2 |
| N304 | Y369 | 4.72 | 5.6 |
| N305 | T368 | 5.44 | 6.2 |
| N305 | Y369 | 5.79 | 6 |
| H312 | Y356 | 5.43 | 17.7 |
| S313 | S356 | 5.96 | 18 |
| S313 | T357 | 5.63 | 16.2 |
| S313 | R358 | 5.92 | 12.9 |
| Y314 | R358 | 4.68 | 12.3 |
| V315 | V359 | 5.19 | 12.2 |
| D316 | I360 | 5.96 | 12.3 |
| S320 | G234 | 5.86 | 25.5 |

**c per residue pLDDT score for AF3 models (BSAP-1, BSAP-1 with OMP-S, and BSAP-1 with OMP-R). Differences in pLDDT for putative complexes normalized to BSAP-1 values (ΔpLDDT) are shown.**

| | pLDDT | | | ΔpLDDT | |
|---|---|---|---|---|---|
| **Residue** | **BSAP-1** | **BSAP-1 + OMP-S** | **BSAP-1 + OMP-R** | **BSAP-1 + OMP-S** | **BSAP-1 + OMP-R** |
| **G338** | 34.37 | 53.57 | 31.16 | 19.20 | -22.41 |
| **G339** | 30.07 | 60.37 | 28.35 | 30.30 | -32.02 |
| **A340** | 31.16 | 61.08 | 27.06 | 29.92 | -34.02 |
| **V341** | 33.76 | 60.29 | 25.66 | 26.53 | -34.63 |
| **R342** | 31.94 | 64.70 | 25.00 | 32.76 | -39.70 |
| **N343** | 32.81 | 57.54 | 24.14 | 24.73 | -33.40 |
| **N344** | 35.73 | 44.34 | 23.81 | 8.61 | -20.53 |
| **S345** | 33.78 | 45.19 | 21.39 | 11.41 | -23.80 |
| **R346** | 31.79 | 43.56 | 23.76 | 11.77 | -19.80 |
| **G347** | 28.18 | 45.69 | 23.35 | 17.51 | -22.34 |
| **H348** | 31.78 | 51.45 | 23.69 | 19.67 | -27.76 |
| **N349** | 25.85 | 59.17 | 23.07 | 33.32 | -36.10 |
| **T350** | 25.14 | 66.30 | 22.72 | 41.16 | -43.58 |
| **S351** | 24.29 | 73.76 | 24.28 | 49.47 | -49.48 |
| **S352** | 22.43 | 69.43 | 25.51 | 47.00 | -43.92 |
| **Y353** | 25.51 | 72.82 | 26.41 | 47.31 | -46.41 |
| **G354** | 23.98 | 67.17 | 28.55 | 43.19 | -38.62 |
| **N355** | 23.94 | 72.74 | 28.63 | 48.80 | -44.11 |
| **S356** | 22.52 | 71.74 | 28.24 | 49.22 | -43.50 |
| **T357** | 21.90 | 69.92 | 30.23 | 48.02 | -39.69 |
| **R358** | 20.66 | 75.57 | 33.39 | 54.91 | -42.18 |
| **V359** | 21.57 | 69.99 | 36.49 | 48.42 | -33.50 |
| **I360** | 20.63 | 72.50 | 35.20 | 51.87 | -37.30 |
| **G361** | 22.41 | 68.14 | 37.32 | 45.73 | -30.82 |
| **A362** | 24.94 | 56.81 | 36.51 | 31.87 | -20.30 |
| **N363** | 21.88 | 52.59 | 36.85 | 30.71 | -15.74 |
| **G364** | 25.34 | 50.44 | 41.81 | 25.10 | -8.63 |
| **N365** | 24.44 | 54.03 | 42.81 | 29.59 | -11.22 |
| **R366** | 22.23 | 54.70 | 50.31 | 32.47 | -4.39 |
| **S367** | 26.07 | 56.67 | 54.05 | 30.60 | -2.62 |
| **T368** | 25.40 | 60.14 | 60.60 | 34.74 | 0.46 |

| | | | | | |
|---|---|---|---|---|---|
| **Y369** | 24.51 | 68.31 | 67.10 | 43.80 | -1.21 |
| **R370** | 28.08 | 67.17 | 69.79 | 39.09 | 2.62 |
| **R371** | 24.69 | 67.97 | 68.47 | 43.28 | 0.50 |
| **P372** | 16.89 | 56.79 | 60.56 | 39.90 | 3.77 |

**Supplementary Table 2. Pixel intensities of the far-western blot measured for peptides comprised of all possible amino acid substitutions normalized to value of the wild type sequence.**

| Peptide array normalised values | | | | | | | | | | | | | | | | | | | | | | | | | | | | | | | | | | |
|---|---|---|---|---|---|---|---|---|---|---|---|---|---|---|---|---|---|---|---|---|---|---|---|---|---|---|---|---|---|---|---|---|---|---|
| | G | A | V | R | N | N | S | R | G | H | N | T | S | S | Y | G | N | S | T | R | V | I | G | A | N | G | N | R | S | T | Y | R | R | P |
| C | 0.954 | 1.111 | 1.532 | 1.093 | 0.929 | 1.028 | 2.233 | 0.996 | 0.967 | 1.039 | 1.037 | 0.920 | 2.045 | 1.913 | 0.740 | 0.790 | 0.912 | 1.239 | 2.270 | 0.360 | 0.523 | 0.447 | 0.375 | 0.512 | 0.432 | 0.436 | 0.436 | 0.574 | 0.345 | 0.444 | 0.157 | 0.934 | 0.383 | 0.756 |
| P | 0.525 | 0.369 | 0.645 | 0.471 | 0.435 | 0.517 | 1.488 | 0.769 | 0.721 | 0.744 | 0.734 | 0.396 | 0.795 | 1.061 | 0.543 | 0.640 | 0.790 | 1.076 | 2.321 | 0.669 | 0.863 | 0.907 | 0.847 | 1.202 | 1.291 | 1.041 | 1.169 | 1.504 | 1.133 | 1.045 | 0.681 | 1.183 | 0.619 | 1.000 |
| G | 1.000 | 1.029 | 1.331 | 0.801 | 0.826 | 0.895 | 1.938 | 0.939 | 1.000 | 0.941 | 1.127 | 1.077 | 2.200 | 2.156 | 1.043 | 1.000 | 1.187 | 1.541 | 2.738 | 0.877 | 1.004 | 0.929 | 1.000 | 1.411 | 1.180 | 1.000 | 1.069 | 1.331 | 0.684 | 0.861 | 0.373 | 1.003 | 0.316 | 0.580 |
| A | 0.989 | 1.000 | 1.420 | 0.937 | 1.100 | 1.029 | 2.341 | 0.890 | 1.045 | 1.064 | 1.108 | 1.008 | 2.153 | 2.137 | 0.921 | 0.799 | 0.900 | 1.152 | 2.049 | 0.573 | 0.721 | 0.653 | 0.741 | 1.000 | 0.940 | 0.780 | 0.631 | 0.751 | 0.598 | 0.907 | 0.378 | 0.959 | 0.527 | 1.062 |
| V | 1.148 | 0.731 | 1.000 | 0.493 | 0.532 | 0.688 | 1.778 | 0.791 | 0.857 | 1.054 | 1.020 | 1.086 | 1.735 | 1.913 | 0.915 | 0.815 | 0.827 | 1.365 | 2.891 | 0.696 | 1.000 | 0.865 | 0.771 | 1.123 | 1.141 | 1.041 | 1.095 | 1.676 | 0.842 | 1.014 | 0.595 | 1.499 | 0.668 | 0.777 |
| L | 0.830 | 0.827 | 1.037 | 0.526 | 0.534 | 0.752 | 1.947 | 0.748 | 0.983 | 1.104 | 1.167 | 1.136 | 1.944 | 1.983 | 0.948 | 0.966 | 1.075 | 1.517 | 3.035 | 1.035 | 1.005 | 0.838 | 0.811 | 1.307 | 1.210 | 1.021 | 1.103 | 1.357 | 0.939 | 1.173 | 0.731 | 1.474 | 0.728 | 0.882 |
| I | 0.871 | 0.910 | 1.216 | 0.617 | 0.657 | 0.702 | 1.870 | 0.863 | 0.970 | 1.157 | 1.175 | 1.157 | 2.017 | 2.016 | 1.088 | 0.894 | 1.085 | 1.562 | 3.460 | 1.027 | 1.275 | 1.000 | 0.847 | 1.344 | 1.300 | 1.114 | 1.115 | 1.574 | 0.996 | 1.106 | 0.575 | 1.447 | 0.628 | 0.768 |
| M | 0.707 | 0.581 | 1.041 | 0.515 | 0.624 | 0.581 | 1.446 | 0.760 | 0.920 | 1.125 | 1.026 | 1.052 | 1.944 | 2.044 | 0.972 | 0.997 | 1.100 | 1.664 | 3.690 | 1.114 | 1.071 | 0.997 | 0.931 | 1.388 | 1.211 | 0.994 | 1.095 | 1.499 | 0.993 | 1.160 | 0.665 | 1.335 | 0.924 | 0.628 |
| F | 0.641 | 0.541 | 1.097 | 0.740 | 0.787 | 0.704 | 1.898 | 0.857 | 0.913 | 1.174 | 1.153 | 0.822 | 1.815 | 2.073 | 0.973 | 0.942 | 1.134 | 1.700 | 3.392 | 1.438 | 1.255 | 1.217 | 0.984 | 1.451 | 1.387 | 1.264 | 1.276 | 1.639 | 1.106 | 1.286 | 0.981 | 1.497 | 0.837 | 1.011 |
| Y | 1.139 | 0.892 | 0.953 | 0.601 | 0.623 | 0.660 | 1.979 | 0.816 | 0.990 | 1.041 | 1.175 | 1.100 | 1.970 | 1.836 | 1.000 | 0.867 | 0.985 | 1.553 | 3.074 | 0.793 | 1.046 | 0.987 | 0.947 | 1.216 | 1.374 | 1.330 | 1.344 | 1.809 | 1.138 | 1.298 | 1.000 | 1.481 | 0.842 | 0.923 |
| W | 0.985 | 0.613 | 0.765 | 0.563 | 0.498 | 0.485 | 1.664 | 0.621 | 0.684 | 0.872 | 0.999 | 0.876 | 1.396 | 1.165 | 0.605 | 0.551 | 0.608 | 0.846 | 1.586 | 0.510 | 0.545 | 0.517 | 0.473 | 1.026 | 0.999 | 0.900 | 0.817 | 0.940 | 0.811 | 0.976 | 0.552 | 1.203 | 0.672 | 0.810 |
| S | 0.504 | 0.377 | 0.534 | 0.354 | 0.346 | 0.395 | 1.000 | 0.405 | 0.454 | 0.554 | 0.547 | 0.451 | 1.000 | 1.000 | 0.403 | 0.492 | 0.557 | 1.000 | 1.681 | 0.474 | 0.528 | 0.642 | 0.838 | 1.279 | 1.102 | 0.984 | 0.987 | 1.439 | 1.000 | 1.029 | 0.455 | 1.521 | 0.765 | 0.627 |
| T | 0.603 | 0.415 | 0.554 | 0.434 | 0.399 | 0.372 | 0.973 | 0.429 | 0.660 | 0.823 | 0.789 | 0.519 | 0.941 | 0.944 | 0.435 | 0.476 | 0.505 | 0.599 | 1.000 | 0.380 | 0.446 | 0.502 | 0.722 | 0.927 | 0.882 | 0.781 | 0.764 | 1.205 | 0.921 | 1.000 | 0.441 | 1.398 | 0.616 | 0.607 |
| N | 1.132 | 1.054 | 1.412 | 1.083 | 1.000 | 1.000 | 2.529 | 0.988 | 1.034 | 1.088 | 1.000 | 1.000 | 1.988 | 1.831 | 0.770 | 0.725 | 1.000 | 1.228 | 2.385 | 0.606 | 0.784 | 0.589 | 0.673 | 0.926 | 1.000 | 0.886 | 1.000 | 1.097 | 0.831 | 0.834 | 0.400 | 1.125 | 0.702 | 0.661 |
| Q | 1.083 | 1.059 | 1.595 | 0.937 | 0.886 | 0.899 | 2.080 | 0.978 | 1.025 | 1.141 | 1.083 | 1.056 | 2.063 | 1.892 | 0.887 | 0.877 | 1.121 | 1.369 | 2.445 | 0.564 | 0.811 | 0.716 | 0.654 | 0.940 | 0.636 | 0.508 | 0.538 | 0.596 | 0.376 | 0.511 | 0.183 | 0.609 | 0.189 | 0.484 |
| D | 1.083 | 0.980 | 1.655 | 1.062 | 0.976 | 0.980 | 2.445 | 0.962 | 0.979 | 1.221 | 1.112 | 1.016 | 1.907 | 1.700 | 0.817 | 0.753 | 0.908 | 1.134 | 2.443 | 0.518 | 0.563 | 0.682 | 0.753 | 0.887 | 0.869 | 0.811 | 0.905 | 0.817 | 0.726 | 0.818 | 0.356 | 1.244 | 0.571 | 1.126 |
| E | 1.065 | 1.164 | 1.432 | 0.885 | 0.732 | 0.950 | 1.943 | 0.921 | 1.013 | 1.118 | 1.162 | 0.996 | 2.128 | 2.102 | 0.999 | 0.944 | 1.206 | 1.577 | 3.175 | 0.906 | 1.053 | 0.900 | 1.006 | 1.439 | 1.053 | 1.029 | 0.948 | 0.753 | 0.712 | 0.941 | 0.216 | 0.781 | 0.311 | 0.873 |
| H | 1.071 | 1.020 | 1.385 | 0.756 | 0.708 | 0.769 | 2.069 | 0.767 | 0.971 | 1.000 | 1.122 | 1.191 | 2.434 | 2.117 | 1.089 | 1.019 | 1.304 | 1.677 | 2.893 | 1.089 | 1.065 | 0.940 | 0.938 | 1.381 | 1.328 | 1.178 | 1.251 | 1.469 | 0.931 | 1.085 | 0.653 | 1.361 | 0.844 | 0.795 |
| K | 0.697 | 0.778 | 1.033 | 0.525 | 0.455 | 0.708 | 1.674 | 0.753 | 0.884 | 0.920 | 0.990 | 0.897 | 1.889 | 1.972 | 0.846 | 0.845 | 0.859 | 1.106 | 2.889 | 0.772 | 0.812 | 0.583 | 0.513 | 1.126 | 0.912 | 0.705 | 0.575 | 0.878 | 0.650 | 0.622 | 0.295 | 0.769 | 0.402 | 0.400 |
| R | 1.156 | 0.998 | 1.305 | 1.000 | 0.985 | 1.070 | 2.321 | 1.000 | 1.094 | 1.246 | 1.094 | 1.122 | 2.171 | 1.845 | 0.725 | 0.637 | 0.856 | 1.031 | 2.490 | 1.000 | 0.836 | 0.715 | 0.741 | 0.939 | 0.980 | 0.790 | 0.893 | 1.000 | 0.680 | 0.832 | 0.336 | 1.000 | 1.000 | 0.619 |

**Supplementary Table 3**. Cryo-EM data collection and model validation statistics.

| | |
|---|---|
| **EMDB** | 58950 |
| **PDB** | 32KD |
| **Data collection** | |
| Microscope | Titan Krios |
| Voltage (keV) | 300 |
| Magnification | 130000 |
| Exposure (e/Å$^2$) | 50 |
| Defocus range (μm) | 0.9 – 3.0 |
| Detector | |
| Pixel size (Å) | 0.931 |
| No. of movies | 8790 |
| **Reconstruction** | |
| Symmetry imposed | C13 |
| No. of particles (start) | 418428 |
| No. of particles (final) | 53700 |
| Resolution (Å) | 2.68 |
| Resolution range (min, 25th percentile, median, 75th percentile, max) | 2.009, 2.845, 3.366, 6.351, 44.084 |
| Map sharpening B factor (Å$^2$) | -86.5 |
| **Model refinement** | |
| Number of residues | 3809 |
| Protein B factor (Å) | 94.93 |
| R.M.S deviations | |
| Bond lengths (Å) | 0.009 |
| Bond angles (°) | 1.764 |
| **Validation** | |
| MolProbity score | 1.97 |
| Clash score | 5.52 |
| Rotamer outliers (%) | 1.95 |
| Cβ outliers (%) | 0.68 |
| Ramachandran | |
| Favoured (%) | 92.78 |
| Allowed (%) | 6.88 |
| Outliers (%) | 0.34 |

**Supplementary Table 4. *Bacteroides fragilis* strains used in this study**

| **Bacterial strains** | **source** |
|---|---|
| *Bacteroides fragilis* 638R Δ*bfubb* (BF638R_3923) | (3) |
| *Bacteroides fragilis* 638R Δ*bfubb* BSAP-1 R38N | this study |
| *Bacteroides fragilis* 638R Δ*bfubb* BSAP-1 P297W | this study |
| *Bacteroides fragilis* 638R Δ*bfubb* BSAP-1 R358W | this study |
| *Bacteroides fragilis* 638R Δ*bfubb* BSAP-1 Y369E | this study |
| *Bacteroides fragilis* 638R Δ*bfubb* BSAP-1 R358W, Y369E | this study |
| *Bacteroides fragilis* 638R Δ*bfubb* BSAP-1 R366E,R370E,R371E | this study |
| *Bacteroides fragilis* 12905 | (4) |
| *Bacteroides fragilis* CL03T12C07 | (5) |
| *Bacteroides fragilis* CL03T12C07 Δ*fpn* (HMPREF1067_01433) | this study |
| *Escherichia coli* BL21/DE3 (His-BSAP-1) | this study |
| *Escherichia coli* BL21/DE3 (His-BSAP-1 R38N) | This study |

**Supplementary Table 5. Primers used in this study**

| | | |
|---|---|---|
| Delete the fragipain gene (Bf638R_2765) from *B. fragilis* 638R. Flanks cloned into pLGB36. | left flank forward | cgaattcctgcagcccggggctggatattccggtcgac |
| | left flank reverse | tatccgcccaagcaagcagcataagcag |
| | right flank forward | gctgcttgcttgggcggataccggttgg |
| | right flank reverse | ggcatagtatcagatgagtgcgcatttaccgaaaagtatccgtc |
| Delete the fragipain gene (HMPREF1067_01433) from *B. fragilis* CL03T12C07. Flanks cloned into pLGB36. | left flank forward | cgaattcctgcagcccggggctggatattccggtcgac |
| | left flank reverse | tatccgcccaagcaagcagcataagcag |
| | right flank forward | gctgcttgcttgggcggataccggttgg |
| | right flank reverse | ggcatagtatcagatgagtgcgcatttaccgaaaagtatccgtc |

**Supplementary References:**

1. Borgini, S., et al., *Identification of receptor-binding domains of Bacteroidales antibacterial pore-forming toxins.* J Biol Chem, 2026. **302**(2): p. 111113.
2. Lomize AL, Todd SC, Pogozheva ID. *Spatial arrangement of proteins in planar and curved membranes by PPM 3.0*. Protein Sci. 2022. **31**(1):209-220.
3. Chatzidaki-Livanis, M., et al., *Gut Symbiont Bacteroides fragilis Secretes a Eukaryotic-Like Ubiquitin Protein That Mediates Intraspecies Antagonism.* mBio, 2017. **8**(6).
4. L. E. Comstock, A. Pantosti, D. L. Kasper, *Genetic diversity of the capsular polysaccharide C biosynthesis region of Bacteroides fragilis*. Infect Immun, 2000. **68**, 6182-6188.
5. N. L. Zitomersky, M. J. Coyne, L. E. Comstock, *Longitudinal analysis of the prevalence, maintenance, and IgA response to species of the order Bacteroidales in the human gut*. Infect Immun, 2011. **79**, 2012-2020.
